# Life cycle process dependencies of positive-sense RNA viruses suggest strategies for inhibiting productive cellular infection

**DOI:** 10.1101/2020.09.19.304576

**Authors:** Harsh Chhajer, Vaseef A. Rizvi, Rahul Roy

## Abstract

Life cycle processes of positive-strand (+)RNA viruses are broadly conserved across families, yet they employ different life cycle strategies to grow in the cell. Using a generalized dynamical model for intracellular (+)ssRNA virus growth, we decipher these life cycle determinants and their dependencies for several viruses and parse the effect of viral mutations and host cell permissivity. We show that Poliovirus employs rapid replication and virus assembly whereas Japanese Encephalitis virus leverages its higher rate of translation and efficient cellular reorganization compared to Hepatitis C virus. Stochastic simulations of the model demonstrate infection extinction if all seeding viral RNA degrade before establishing robust replication. The probability of productive cellular infection is affected by virus-host processes, defined by early life cycle events and viral seeding. Synergy among these parameters in limiting infection suggests new avenues for inhibiting viral infections by targeting early life cycle bottlenecks.

## Introduction

Positive sense single stranded RNA ((+)ssRNA) viruses, that include *Enteroviridae* (Poliovirus), *Flaviviridae* (Dengue virus), *Coronaviridae* (SARS) virus families, are a major public health challenge. Better understanding of their growth dynamics in the cell can help us identify new drug targets and novel antiviral approaches. Viruses infect and grow inside the cell using a complex set of molecular and cellular processes that has evolved to ensure successful propagation. (+)RNA viruses sequentially translate viral proteins using the positive strand RNA genome upon cell entry, replicate to form nascent genomes from a double stranded RNA (dsRNA) replication intermediate and create new virus particles by encapsidating the (+)RNA genomes with its structural proteins. Members of this class show significant diversity in genome size, physical makeup, constituent viral proteins, host tropism, and chronicity of infection. Yet, they also display striking similarities in cellular life cycle dynamics, closely imitating mechanisms of replication, translation, virus assembly as well as similar interactions with the host cell machinery. This has motivated the search for universal features that can be exploited as broad spectrum anti-viral targets.

One common characteristic of most (+)RNA viruses is the induction of significant alterations of the intracellular host membranes [1, 2, 3, 4]. The vesicular membranous structures formed, protect viral RNA and proteins from cytosolic degradation and host defense systems, provide a conducive micro-environment for efficient viral replication and hence are also referred to as replication compartments (CMs). Impeding membrane re-organization has been shown to decelerate cellular infection dynamics [5, 6], lower viral yield [7, 8, 9, 10] and reduce the propensity of the virus to establish productive cellular infection in the host cell [6, 11, 12]. In general, failure to establish viral infection has been associated with cellular heterogeneity and is attributed to the random loss of genome segments by RNA degradation in the early stages of infection [13, 14, 15]. This stochastic effect is more pronounced at low multiplicity of infection [12] and is observed for many virus families. While this suggests that early events in the virus life cycle define the fate of infection, what factors control this mode of viral clearance have not been profiled in detail.

Quantitative measurements and mathematical modeling has tremendously enhanced our understanding of how the subtleties of intracellular processes shape the outcome during viral infections [16, 17, 18, 19, 20, 21, 22, 23, 24, 25, 26]. Understanding derived from these (and their extensions that incorporate extracellular and immune response) have been used to determine effectiveness of interventions and combinations thereof [27, 28, 29, 30, 31]. However, detailed models with explicit molecular details suffer from redundancy in fitting parameters or challenges with estimating parameters values experimentally. On the other hand, generalized models fail to accurately emulate the experimentally observed dynamics across a large class of viruses, viral strains and different host cells. Nevertheless, viral dynamics models that universally capture experimental observations while allowing sufficient inference of molecular mechanisms across many viruses can be insightful. Apart from identifying life cycle bottlenecks, they can be employed to predict the effectiveness of broadly applicable anti-virals.

Most intracellular +RNA viral dynamics models also do not account for the slow formation kinetics of the membranous replication compartments. Since this event coincides with the early infection that is known to be sensitive to stochastic fluctuations in viral RNA, membrane reorganization can influence the virus life cycle in complex ways. In this study, we extend previous viral dynamics models [19, 17, 22] by incorporating the kinetics of replication compartment formation post infection. We show that our model accurately captures several variants of experimentally measured dynamics for Hepatitis C virus (HCV)[22], Japanese Encephalitis virus (JEV) [32] and Poliovirus (PV) [19] infection from literature. Further, it identifies differences among virus life traits and dynamics and describes effect of viral mutations[5] and host factor silencing [7, 5]. We show that the dynamics of compartment formation is a critical kinetic bottleneck for the viruses. It influences the ability of various viruses to establish a productive infection in the host cell that we refer here as ‘cellular infectivity’ (Φ). Apart from replication, we recognize limiting formation of CM, increased cytoplasmic degradation of viral RNA and reduced translation as pan-viral strategies, and estimate the synergy among them in limiting cellular infectivity.

## Results

### Cellular viral life cycle model for monopartite (+)RNA viruses

We propose a mathematical model for cellular life cycle of single stranded monopartite (+)RNA viruses (Fig 1a, details in SI S1). It is inspired by previous HCV and PV models [19, 17, 22] but we focus on intracellular processes common to such viruses and lump molecular details when appropriate (eq. 2, Table 1). Upon entry into the cell, viral (+)RNA in cytoplasm (*R*_*cyt*_) is translated by host ribosomes to produce structural (*P*_*S*_) and non-structural (*P*_*NS*_) proteins at a rate *k*_*t*_ (eqs. 2.2, 2.3). Though translation occurs in the cytoplasm, replication is mainly restricted to replication compartments (CMs) [1, 2].

**Table 1:**
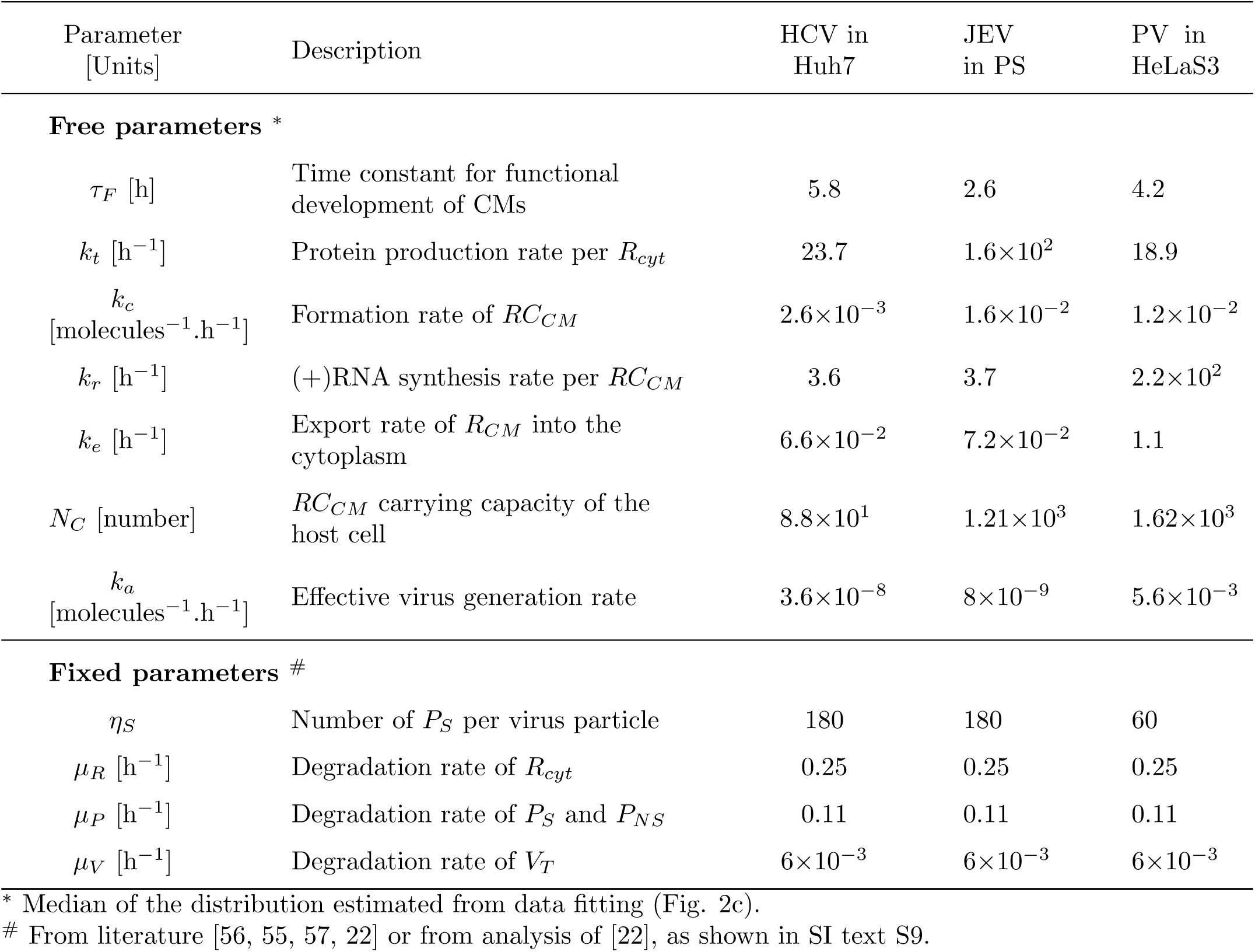
Life cycle Model Parameters.

| Parameter<br>[Units] | Description | HCV in<br>Huh7 | JEV<br>in PS | PV in<br>HeLaS3 |
| --- | --- | --- | --- | --- |
| **Free parameters** * |  |  |  |  |
| $\tau_F$ [h] | Time constant for functional development of CMs | 5.8 | 2.6 | 4.2 |
| $k_t$ [h<sup>-1</sup>] | Protein production rate per $R_{cyt}$ | 23.7 | $1.6 \times 10^2$ | 18.9 |
| $k_c$<br>[molecules<sup>-1</sup>.h<sup>-1</sup>] | Formation rate of $RC_{CM}$ | $2.6 \times 10^{-3}$ | $1.6 \times 10^{-2}$ | $1.2 \times 10^{-2}$ |
| $k_r$ [h<sup>-1</sup>] | (+)RNA synthesis rate per $RC_{CM}$ | 3.6 | 3.7 | $2.2 \times 10^2$ |
| $k_e$ [h<sup>-1</sup>] | Export rate of $R_{CM}$ into the cytoplasm | $6.6 \times 10^{-2}$ | $7.2 \times 10^{-2}$ | 1.1 |
| $N_C$ [number] | $RC_{CM}$ carrying capacity of the host cell | $8.8 \times 10^1$ | $1.21 \times 10^3$ | $1.62 \times 10^3$ |
| $k_a$<br>[molecules<sup>-1</sup>.h<sup>-1</sup>] | Effective virus generation rate | $3.6 \times 10^{-8}$ | $8 \times 10^{-9}$ | $5.6 \times 10^{-3}$ |
| **Fixed parameters** # |  |  |  |  |
| $\eta_S$ | Number of $P_S$ per virus particle | 180 | 180 | 60 |
| $\mu_R$ [h<sup>-1</sup>] | Degradation rate of $R_{cyt}$ | 0.25 | 0.25 | 0.25 |
| $\mu_P$ [h<sup>-1</sup>] | Degradation rate of $P_S$ and $P_{NS}$ | 0.11 | 0.11 | 0.11 |
| $\mu_V$ [h<sup>-1</sup>] | Degradation rate of $V_T$ | $6 \times 10^{-3}$ | $6 \times 10^{-3}$ | $6 \times 10^{-3}$ |

\* Median of the distribution estimated from data fitting (Fig. 2c).

**Figure 1:**
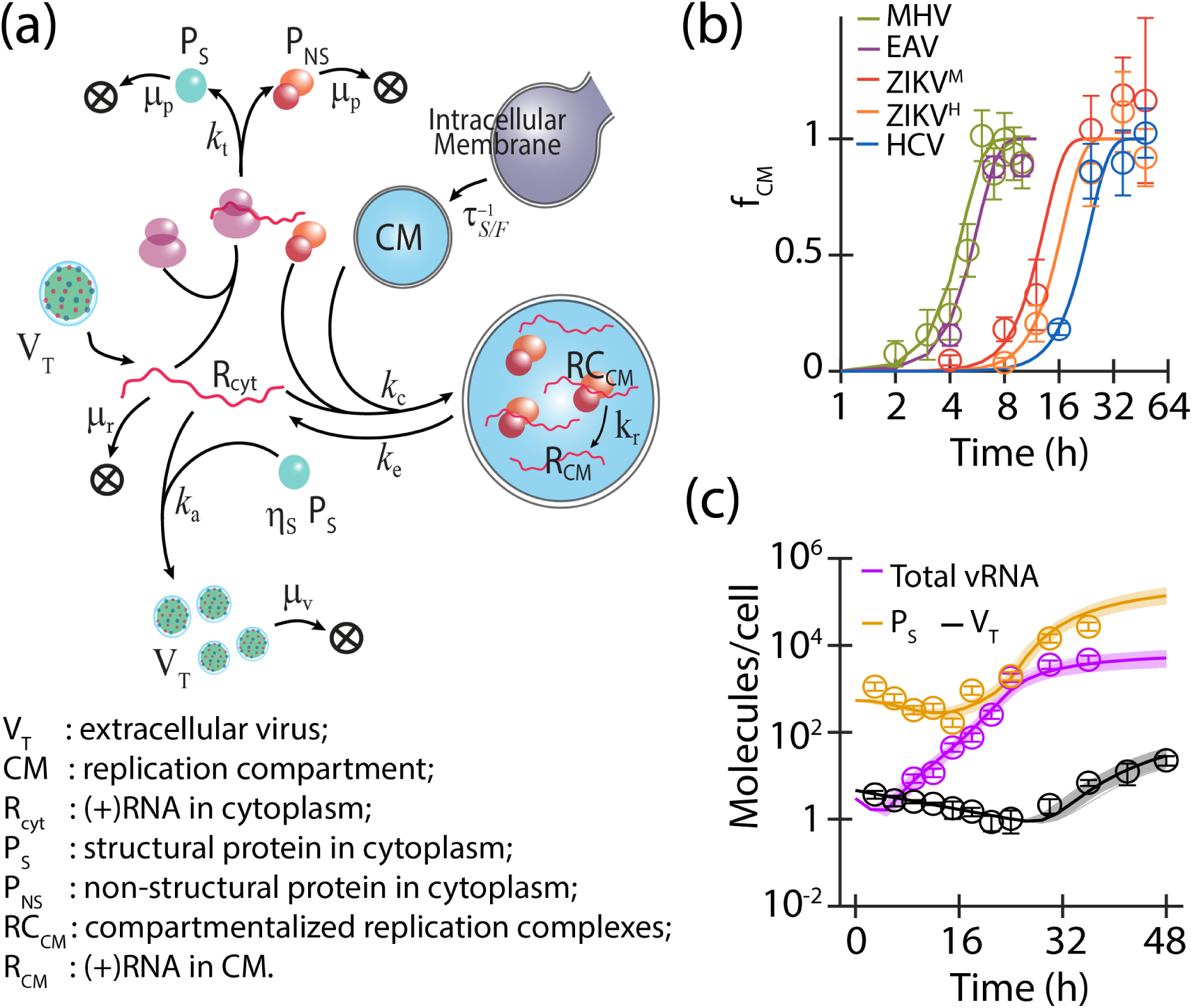
Viral life cycle model and incorporation of compartment formation dynamics. (a) Schematic of the viral life cycle model. In the cytoplasm, the (+)RNA (*R*_*cyt*_) is translated by the host ribosomes to produce viral structural (*P*_*S*_) and non-structural proteins (*P*_*NS*_). *η*_*S*_ copies of *P*_*S*_ associates with *R*_*cyt*_ to assemble virus particles (*V*_*T*_). Intracellular membrane is re-organized to form compartments (CM) which harbours viral replication complex (*RC*_*CM*_) that produce new (+)RNA strands, which are exported out into the cytoplasm. (b) Normalized dynamics of compartment formation (*f*_*CM*_) observed for different (+)RNA viruses fit (lines) using eq. 1 (n = 4) is shown. Data is derived from [37, 38, 39, 40] for Mouse hepatitis virus (MHV), Equine arteritis virus (EAV), MR766 and H/PF/2013 strains of Zika virus (ZIKV^*M*^ and ZIKV^*H*^) and Hepatitis C virus (HCV). (c) Virus life cycle model fit for HCV infection in Huh7 cells [22]. Circles and error bars correspond to data and colored lines represent respective fits. Thin lines (lightly colored in background) represent dynamics predicted using a set of best parameter combinations (250 sets) from iABC and thick lines denotes their average.

CM formation occurs via extensive alteration to intracellular host membranes [33, 34] induced by viral and host proteins post infection [1, 2, 3, 35]. Although a slow and critical step conserved across many (+)RNA viral life cycle, previous models did not account for their gradual formation. We model this dynamics using a functional form based on analysis of cellular ultra-structural characterization (Fig 1b and Figure S1). Assuming that host membrane homeostasis, we use the popular Weibull function [36] to model the normalized growth of replication compartments, *f*_*CM*_ (eq. 1).

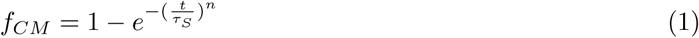

Here, *τ*_*S*_ parameterizes the time scale of the structural manifestation of CMs whereas *n* defines the steepness of the change. Fig 1b shows that increase in vesicular membranous structures observed among (+)RNA viruses is indeed virus and strain specific [37, 38, 39, 40]. Though the value of *τ*_*S*_ does not vary significantly, fitting improves as n increases from 2 to 4, suggesting synchrony in CM generation (Table S1). *τ*_*S*_ estimates generally correlate with the timescale of cellular infection across viruses and strains as observed for the Zika strains [41].

In context of viral replication, one must consider the ability of these sites to provide protective confinement for the RNA replication complex but this aspect has not been quantitatively characterized. Since the structural and functional aspects are likely to be correlated, we use the same functional form (eq. 1 with n=4), but consider a different maturation time (*τ*_*F*_) to model the functional maturation of CMs for replication hereon. Although not explicitly incorporated, *τ*_*F*_ subsumes other delays associated with virus entry and genome un-coating as well. However, such delays have been shown to be comparatively smaller [42, 19, 22].

The number of available CMs limit the replication complex formation. *R*_*cyt*_ and *P*_*NS*_, compete for the unoccupied compartments given by 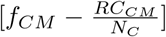, where *RC*_*CM*_ denotes the number of compartmentalized replication complexes and *N*_*C*_ is the carrying capacity associated with *RC*_*CM*_ (Table 1). Therefore, we model the compartmentalization of replication with a logistic function (eqs. 2.1, 2.4). *RC*_*CM*_ synthesize new (+)RNA strands at a rate, *k*_*r*_ (eq. 2.4). The (+)RNA in the compartments (*R*_*CM*_) are exported out into the cytoplasm at rate, *k*_*e*_ (eqs. 2.1, 2.5), where it can re-participate in the life cycle. While the viral RNA and proteins degrade in the cytoplasm, we ignore degradation in the compartments [4, 2, 26]. Finally, viral assembly occurs in cytoplasm where *R*_*cyt*_ associates with *η*_*S*_ molecules of *P*_*S*_ to produce extracellular viral particles (*V*_*T*_) at an overall rate of *k*_*a*_ (eqs. 2.3, 2.6).

### Recapitulating observed HCV life cycle dynamics

Using an iterative approximate Bayesian approach (iABC, see Methods and SI SM1), we fit our viral life cycle model to the dynamics of intracellular viral RNA, proteins, and extracellular viruses observed and recover parameter estimates for the well characterized HCV infection in Huh7 cells [22] (Fig 1c, Tables 1, S2). We find that new (+)RNA strands are produced at (*k*_*r*_=) 3.6 *h*^−1^ per compartmentalized replication complex (*RC*_*CM*_). Using RNA polymerization rate of 150 *nt/min* [43], *we predict around three simultaneous replication events per RC*_*CM*_. This is consistent with experimental observations of synthesis of multiple viral RNA per replication intermediate (reported to be 5 for the closely related Dengue virus [44]). Similarly, our steady state ratio of viral (+)RNA to (-)RNA (= 54:1) compares well with the experimentally observed ratio of 30:1 [45, 46].

While differences in parameter definition limit exact comparison, we find good agreement with previous efforts to model HCV dynamics. For example, approximating 10 ribosomes [47] translating the viral RNA at a time and *k*_*t*_ = 23.7 *h*^−1^, we predict the HCV protein production rate to be 2.4 *h*^−1^ per RNA. This is comparable to the previous estimate of the rate limiting step in protein synthesis, the polyprotein cleavage rate at 1 *h*^−1^ [16, 17]. Our prediction for (+)RNA export out of replication compartments is also similar to previous estimates [16]. However, virus production in our model is 50-fold faster than previous estimates. We attribute this to the unaccounted delay in CM formation that is neglected in prior models and this contributes to the reduced effective assembly rates [22]. Overall, we recapitulate HCV experimental observations not built a priori into the model as well as match previous estimates for comparable parameters.

To further validate our model, we evaluate the life cycle dynamics of subgenomic HCV (sgHCV) transfected into Huh7 (Huh7-Lp) and it highly permissive derivative (Huh7-Lunet) cells using our model [17] (Figure S2). Our estimates for *k*_*t*_, *k*_*r*_ and *k*_*c*_ (the rate of formation of *RC*_*CM*_) for the subgenomic viral transfection are similar to corresponding estimates for full-genomic HCV infection (Table S2, Table S3) suggesting robustness of our model across different experimental systems for HCV. However, the sgHCV system exhibits delayed RC formation and faster (+)RNA export out of CM likely due to lack of structural proteins, transfection induced cellular artifacts or additional pre-processing required for transfected RNA [17]. Importantly, the highly permissive cell line (Huh7-Lunet) exhibits faster CM formation (1.9 fold lower *τ*_*F*_) and higher stability of viral dsRNA replication intermediate (11.8 fold larger *N*_*C*_) compared to Huh7-Lp [48], suggesting efficient replication compartmentalization contributes to higher cellular permissivity (Table S3).

### Comparative analysis of monopartite (+)RNA viruses

To understand the differences in life cycle traits among (+)RNA viruses, we further evaluate our model with two distinct families of viruses for which comprehensive viral dynamics data exists, namely *Enteroviridae* (PV [19]) and *Flaviviridae* (JEV [32]). Comparison of life cycle process parameters (Fig 2, Table S2) shows that the replication rate (*k*_*r*_) and export rate of (+)RNA from compartment (*k*_*e*_) exhibit virus family specific trends (Fig 2c). For example, Poliovirus RNA replicates rapidly (60-fold higher *k*_*r*_) and re-enters the cytoplasmic pool faster (16-fold higher *k*_*e*_) than *Flaviviridae* family (HCV and JEV). Using our estimate of *k*_*r*_ = 210 *h*^−1^ for PV (similar to 133 *h*^−1^ estimated earlier [19]) and assuming PV genome replication takes 100 secs [49], we predict 5.8 simultaneous replication events occur per *RC*_*CM*_ closely matching previously measured values of 6.5 −7 [50, 51]. On the other hand, JEV displays ≈ 3 simultaneous replications per *RC*_*CM*_ (assuming RNA elongation rate of 150 *nt/min* [43]) comparable to HCV and Dengue. This further shows that our model captures the now well recognized fact that CMs are sites of multiple parallel replication reactions without explicitly assuming it [2].

**Figure 2:**
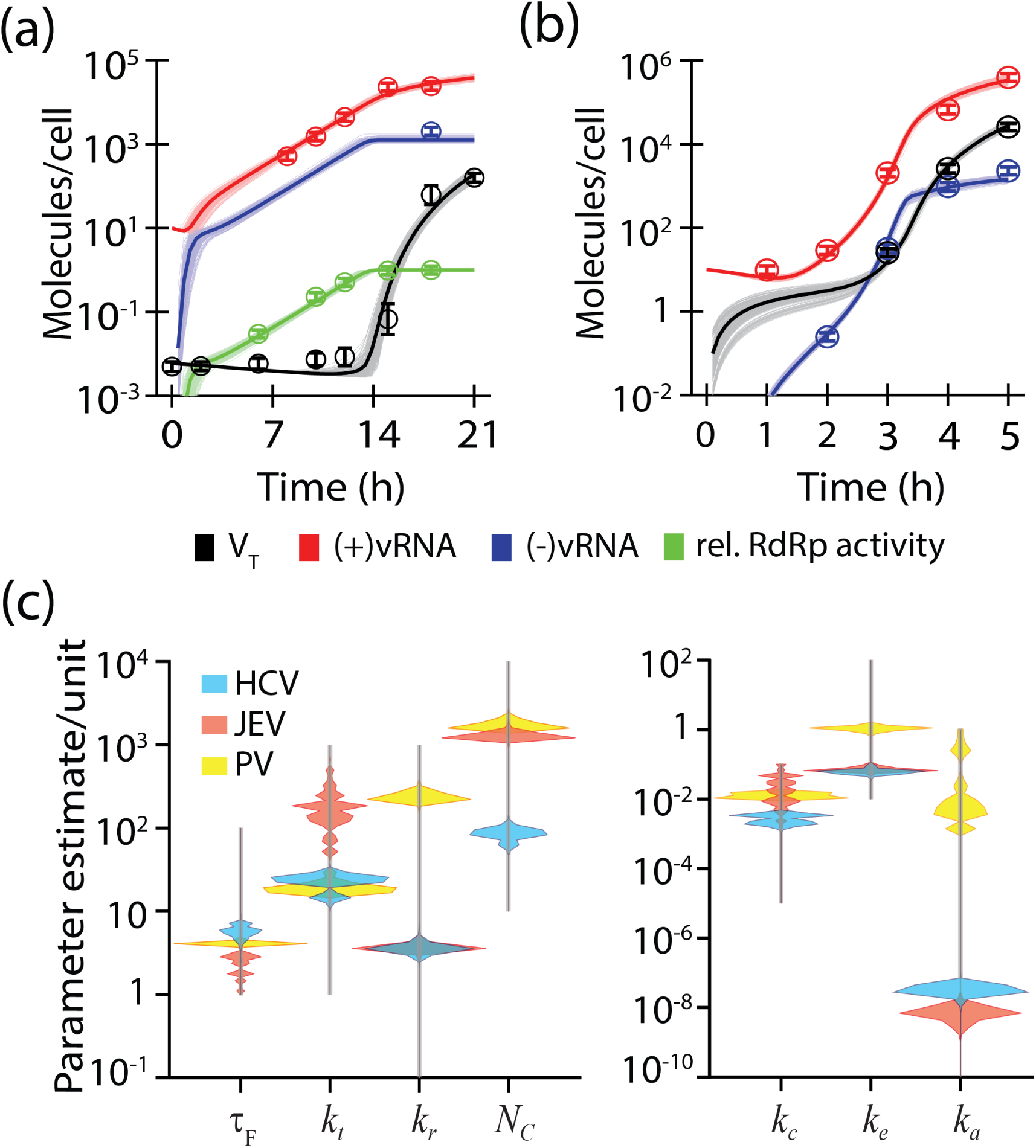
Comparison of monopartite (+)RNA viruses. Model fits (thin lines, average shown as a thick line) for the intracellular dynamics of virus constituents in case of (a) JEV infection in PS cells [32], and (b) PV infection in HeLaS3 cells [19] are shown. (c) Comparison of parameter value distributions estimated for HCV, JEV and PV life cycle dynamics from model fitting. Grey vertical lines indicate the range of initial guesses used for each parameter.

Protein synthesis rate, *k*_*t*_ = 18.9 *h*^−1^ for PV is comparable to HCV and similar to previous report [19] but JEV proteins are produced seven times faster. Although polyprotein processing and host cell state affect *k*_*t*_, we attribute the high *k*_*t*_ values for JEV to its RNA cap dependent translation initiation [52] compared to the IRES mediated mechanism employed by HCV and PV [53, 54]. Faster protein production and an associated early induction of membrane re-organization could contribute to the faster functional maturation of CM for JEV consistent with our estimates for *τ*_*F*_.

Virus assembly and generation defined by *k*_*a*_ is significantly (*>* 10^5^ fold) faster for PV compared to the *Flaviviridae* viruses (HCV and JEV) reflecting their corresponding complexity in assembly and maturation. While the detailed mechanisms of virus assembly remains poorly understood, HCV and JEV are enveloped viruses made of 180 copies of three different structural proteins [55, 56] that require maturation post assembly whereas PV is a smaller non-enveloped virus [57].

These differences in virus life cycle traits are robust to alternative formulations (SI S5). For example, whether we consider pre-formed CM (*f*_*CM*_ = 1) [Figure S5], or a more gradual rise in compartments (Weibull exponent, n = 2) for replication compartment formation [Figure S6], high rates for JEV translation, and rapid replication and assembly of PV sets them apart (Table S4). Similarly, stoichiometry of *P*_*NS*_ in formation of *RC*_*CM*_ [Figure S7], or consideration of replication coupled assembly of viral particles [22] [Figure S8] does not alter our virus specific parameter trends. While goodness of fit based on cumulative AIC values for the three viruses (Table S4) demonstrates a marginal advantage in favour of our main model, measured CM formation dynamics [33, 34] lend additional support to our choice of the model. Similarly, independent experimental data like recovery of steady state levels of replication intermediate [58, 17, 59] and steady state positive/negative RNA ratios [46, 45] are congruent with our model (Table S4). Due to the lack of molecular details for virus assembly, our model only qualitatively captures the virus assembly and particle release dynamics and we cannot discriminate between alternate sites (CMs vs cytoplasm) for virus assembly.

### Conserved and virus-specific determinants shaping viral life cycle

To evaluate how perturbations in life cycle model parameters affect the viral dynamics, we employed temporal sensitivity analysis (TSA) using the eFAST algorithm ([60]). TSA profiles of *RC*_*CM*_, the key intermediate and a surrogate for viral replication, highlights three distinct phases for the viruses (Fig 3 a). The initial establishment (E) phase is sensitive to the delay in formation of CM (*τ*_*F*_), and displays minimal replication due to shortage of CM. The next growth (G) phase represents the rapid increase in viral RNA production and is influenced by parameters governing the increase of (+)RNA in the cytoplasm, and thus the formation of dsRNA replication intermediate. Growth phase is sensitive to changes in viral replication rate (*k*_*r*_), the kinetics of (+)RNA export from CM (*k*_*e*_) and the rate of cytosolic degradation of (+)RNA (*µ*_*R*_). The final saturation (S) phase is defined by the pseudo-steady state behaviour primarily regulated by the carrying capacity for *RC*_*CM*_ (*N*_*C*_). Though the TSA trends are qualitatively similar, the time associated with each phase varies with the virus. The length of the E phase correlates with the estimate for *τ*_*F*_, and time span of growth phase reflects 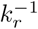 and 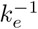. So, while the G phase is comparable for HCV and JEV, it is very short for PV as is evident with the rapid increase in PV RNA in a short window of 2 h [19].

**Figure 3:**
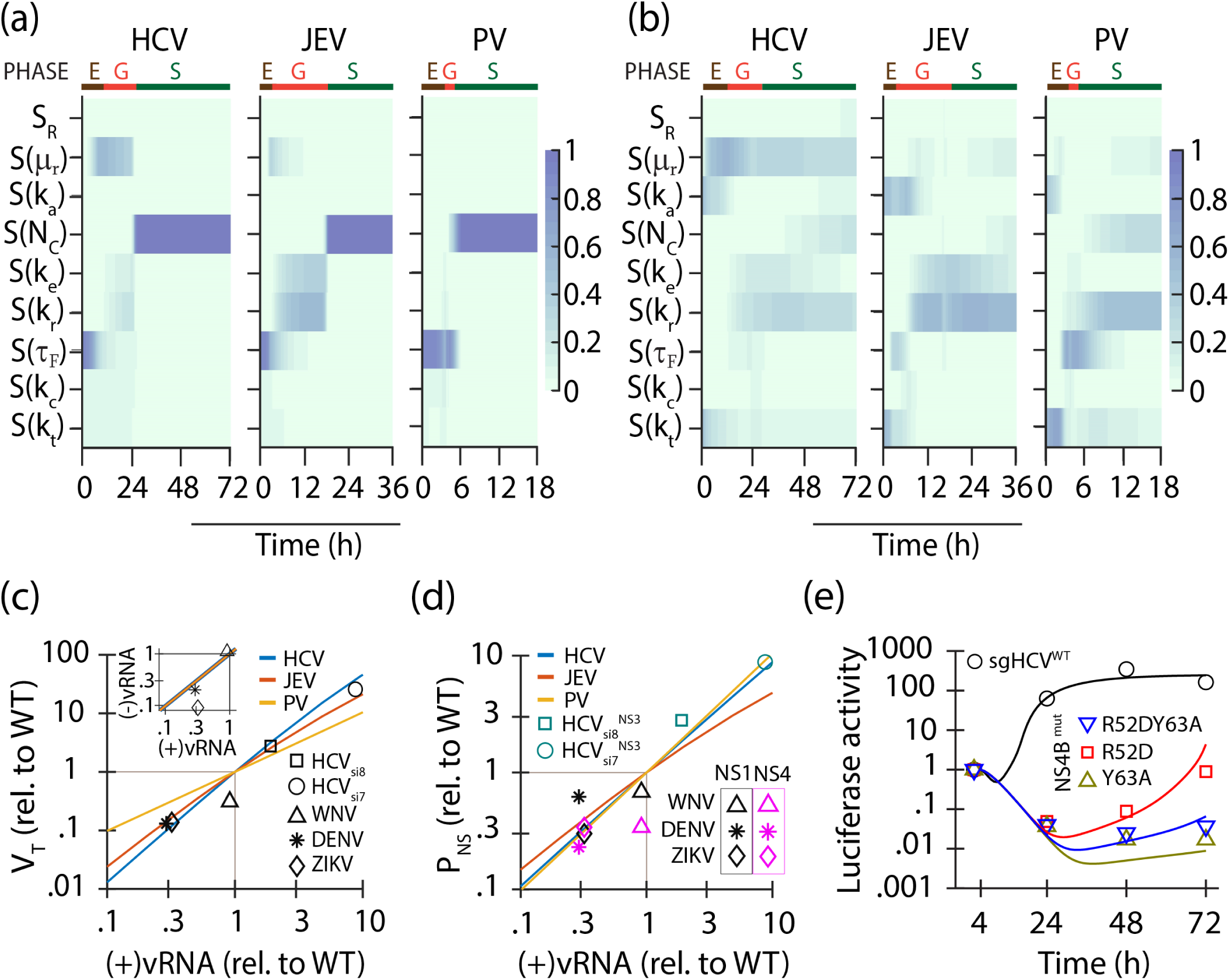
Perturbation analysis of viral life cycle parameters. Parameter-temporal sensitivity profiles for dynamics of (a) *RC*_*CM*_ and (b) *V*_*T*_, for HCV, JEV and PV (with initial seeding of *R*_*cyt*_=3) are shown. *S*(*X*) denotes the profile associated with parameter, X, and *S*_*R*_ = *S*(*µ*_*P*_) + *S*(*µ*_*V*_) + *S*(*dummy*). Based on the *RC*_*CM*_ profiles, life cycle of each of the viruses can be divided into the establishment (E), the growth (G) and the saturation (S) phases as shown. Time axis is not to scale across profiles for different viruses. Fold change in steady state levels (c) *V*_*T*_ vs (+)RNA, (d) *P*_*S*_ vs (+)RNA and (c-inset) (-)RNA vs (+)RNA, due to change in *N*_*C*_ is compared to experimentally data [7, 5]. Solid lines are model predictions and symbols show experimental data. (e) Viral protein dynamics for NS4B mutants is fit with life cycle model by varying *τ*_*F*_. Solid lines and symbols show best fits and data [5], respectively.

Differences in the TSA profiles across the viruses are more evident when *V*_*T*_ (Fig 3b) and *R*_*cyt*_ are considered (Figure S9). The profiles associated with *V*_*T*_ are particularly informative in identifying ‘choke points’ and their effectiveness for different viruses. We postulate that perturbations to replication (*k*_*r*_) would be more effective than translation (*k*_*t*_) against JEV but vice versa for inhibiting Poliovirus growth. For HCV, viral dynamics is influenced by viral RNA degradation (*µ*_*R*_) to a large extent followed by translation and replication. *µ*_*R*_ is critical for HCV life cycle as its (+)RNA has a large dwell time in the cytoplasm (due to its large *τ*_*F*_, 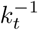 and 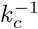). This shows how parameter sensitivity in our model can generate virus-specific insights into intervention strategies.

### Changes in *N*_*C*_ and *τ*_*F*_ mimic the effect of perturbations to compartment formation

Among the life cycle parameters, *τ*_*F*_, *k*_*e*_ and *N*_*C*_ are reflective of the CM formation dynamics and CM morphology. Various viral [1, 2, 4, 35, 61] and host perturbations [62, 63, 64, 65] and drug interventions [9, 8, 10] have been reported to alter membrane reorganization, which ultimately affects infection kinetics as well as the steady state achieved in the late stages of infection. We emulate some of these perturbations by varying *τ*_*F*_ (for kinetics) and *N*_*C*_ (for steady state) and compare it to experimental observations.

Reticulon 3 (RTN3) is an Endoplasmic Reticulum (ER) shaping host protein shown to be involved in ER membrane re-organization during various (+)RNA viral infections [62, 63]. Silencing RTN3 in host cells reduces viral replication of Flaviviruses [7] and Enteroviruses [6], but not in the case of HCV [5]. In our model, *N*_*C*_ is the sole parameter that affects the steady state levels of viral (-)RNA levels (*RC*_*CM*_), which is perturbed upon RTN3 silencing in host cells [7]. By just varying *N*_*C*_, we are able to reproduce correlated fold changes in (-)RNA levels, viral titre and *P*_*NS*_ with respect to viral RNA as observed for various Flaviviruses[7] and HCV[5] upon silencing RTN3 (Fig. 3c-inset, c, d). This also suggests that the steady state level correlations among the various virus constituents are appropriately captured by our model.

Our steady state level relations for levels of virus and viral protein with viral (+)RNA are distinct for the three viruses considered here. For HCV and PV, *P*_*NS*_ varies linearly with viral (+)RNA level, however it is sub-linear in case of JEV (Fig. 3d). The increase in *V*_*T*_ with viral (+)RNA levels is super-linear and sub-quadratic, for JEV and HCV, respectively whereas it is linear for PV (Fig. 3d). The trends are corroborated by steady state analysis of the model (refer SI S3). Efficient assembly for PV (*k*_*a*_*R*_*cyt*_ *>> µ*_*P*_) leads to *R*_*cyt*_-independent level of *P*_*S*_ 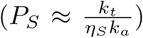, resulting in a linear relation between *V*_*T*_ and 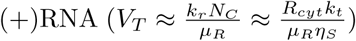. When comparing HCV and JEV, higher *k*_*t*_.*k*_*a*_ estimate contributes to faster assembly of *R*_*cyt*_. Thus, *R*_*cyt*_ (and consequently 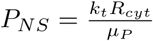) increases sub-linearly with *N*_*C*_ for JEV.

To evaluate the effect of the compartment formation kinetics, we compare viral polyprotein dynamics of HCV NS4B mutants, shown to be defective in inducing membrane re-organization [35, 61]. Using *τ*_*F*_ as the sole fitting parameter (details in SI SM3), the model is able to accurately recover the normalized protein dynamics observed for these sgHCV mutants [5] (Fig 3e). Large *τ*_*F*_ estimated for the NS4B sgHCV mutants R52D, Y63A and R52DY63A, (63, 101 and 80 h, respectively) compared to 5.8 h for the WT virus highlights how increased delay in CM formation affects viral dynamics.

### Compartmentalization of replication defines the fate of virus infection

Compartmentalization of viral replication establishes sites for efficient (+)RNA replication, protected from cytoplasmic degradation in the infected cell. However, compartmentalization is not always guaranteed upon virus entry, with the possibility of degradation of viral genome in the host cytoplasm before membrane reorganization. This aspect was not considered in models that assume pre-existing CMs. We posit that the infection outcome of viral seeding event is an all-or-none phenomena that is determined at the onset of the infection by the opposing effects of cytoplasmic viral RNA degradation and the formation of *RC*_*CM*_ (Fig. 4a). Indeed, stochastic simulations of the HCV life cycle demonstrates these two outcomes similar to previous reports [26] (Fig. 4b). All realizations where *RC*_*CM*_ is formed before the complete degradation of viral RNA result in a productive infection, otherwise the infection extinguishes. We define this likelihood of productive infection establishment post virus seeding as ‘cellular infectivity’, Φ.

**Figure 4:**
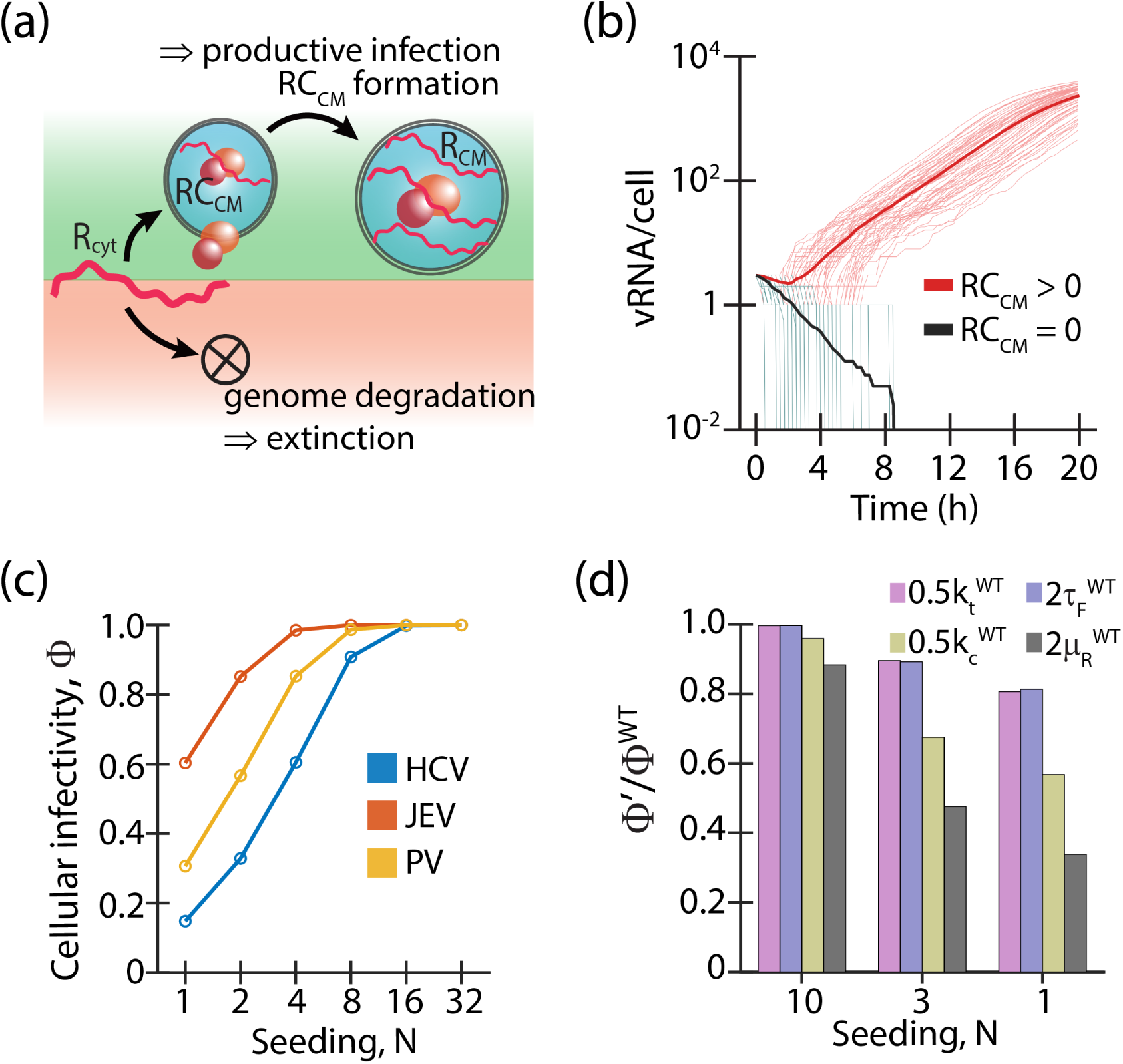
Stochastic fate of viral infection. (a) When the cytoplasmic viral RNA is encapsulated in vesicular compartments, high replication rate and minimal degradation leads to productive infection. Otherwise, degradation of all viral RNA in the cytoplasm leads to complete extinction of the infection. (b) Stochastic realizations of cellular life cycle dynamics of HCV in Huh7 cells at viral seeding of 3. Thin lines show simulated trajectories for viral RNA, (red) when at least 1 *RC*_*CM*_ was formed or (black) when no *RC*_*CM*_ were formed and thick lines represent their corresponding averages. (c) Probability of establishment of infection (Φ) as a function of seeding number (N) for HCV, JEV and PV is shown. (d) Fold change in Φ for PV infection corresponding to a change in various parameters with respect to the wild type (WT, Table 1) values at different virus seeding (N) is shown.

Φ ranges between zero (deterministic extinction of infection) and one (deterministic establishment of productive infection), and is influenced by virus-host factors that are critical in the early stages of virus life cycle. For example, Φ increases monotonically to saturation with viral seeding load (N) (Fig 4c). It is also modulated by the kinetics of viral processes leading to compartmentalization of replication (*k*_*t*_, 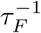, *k*_*c*_) and the stability of viral genome in host cytoplasm 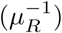 [Fig 4d, Figure S10, Figure S11]. This is consistent with reports that show reduction in infection success rate by Phosphatidylinositol 4-kinase III alpha silencing [11] or for Enterovirus mutants [6] where CM formation is expected to be hindered. This effect is also associated with the antiviral activity of membrane re-modeling inhibitors like K22 [9, 8] that likely influence *τ*_*F*_. Φ also increases from HCV to PV to JEV, based on our estimates of their life cycle parameters (Fig 4c). We estimate a significant fraction of single virus infections are non-productive (estimated to be 40% and 85% for JEV and HCV, respectively). With larger viral seeding, Φ increases such that at N=8, Φ of 0.95 closely matches the 95.28% infection success rate observed with HCV (8 RNA measured inside the cell at 3 hpi [22]).

Interestingly, while infection success rate with 10 seeding genomes remains unaffected when *τ*_*F*_ is increased, it drops by 47% for single genome infection for the same (by 2-fold) change in *τ*_*F*_. This mirrors the larger reduction in fraction of productive PV infected cells observed due to action of membrane re-organization inhibitor, PIK93 [66] at low multiplicity of infection [12]. This effect also contributes to the synergy observed between entry inhibitors (that would decrease effective N) and other antiviral agents, like protease inhibitors, membrane re-organization inhibitors, cyclophilin inhibitors, against HCV [67, 68].

### Synergy among strategies reducing cellular infectivity

Since the life cycle parameters that limit Φ collaborate in complex ways, we conjectured their inter-dependence would give rise to synergy in their action. We use the Bliss independence criterion [69] to evaluate this synergy (Ψ) since these early life cycle events are likely to occur independently at the molecular level. Apart from reducing Φ independently, *τ*_*F*_ and *µ*_*R*_ positively synergize 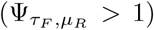 when combined for the viruses (Figs. 5a, Figure S12a,b). For example, doubling of both *τ*_*F*_ and *µ*_*R*_ values resulted in an eight fold reduction in Φ compared to the product of their independent actions in case of HCV. In our formalism, synergy stems from enhanced delays in the formation of compartments leading to increased exposure of viral RNA to cytosolic degradation. By extension, other RNA viruses that employ compartmentalization to stabilize replication will also display such behaviour.

**Figure 5:**
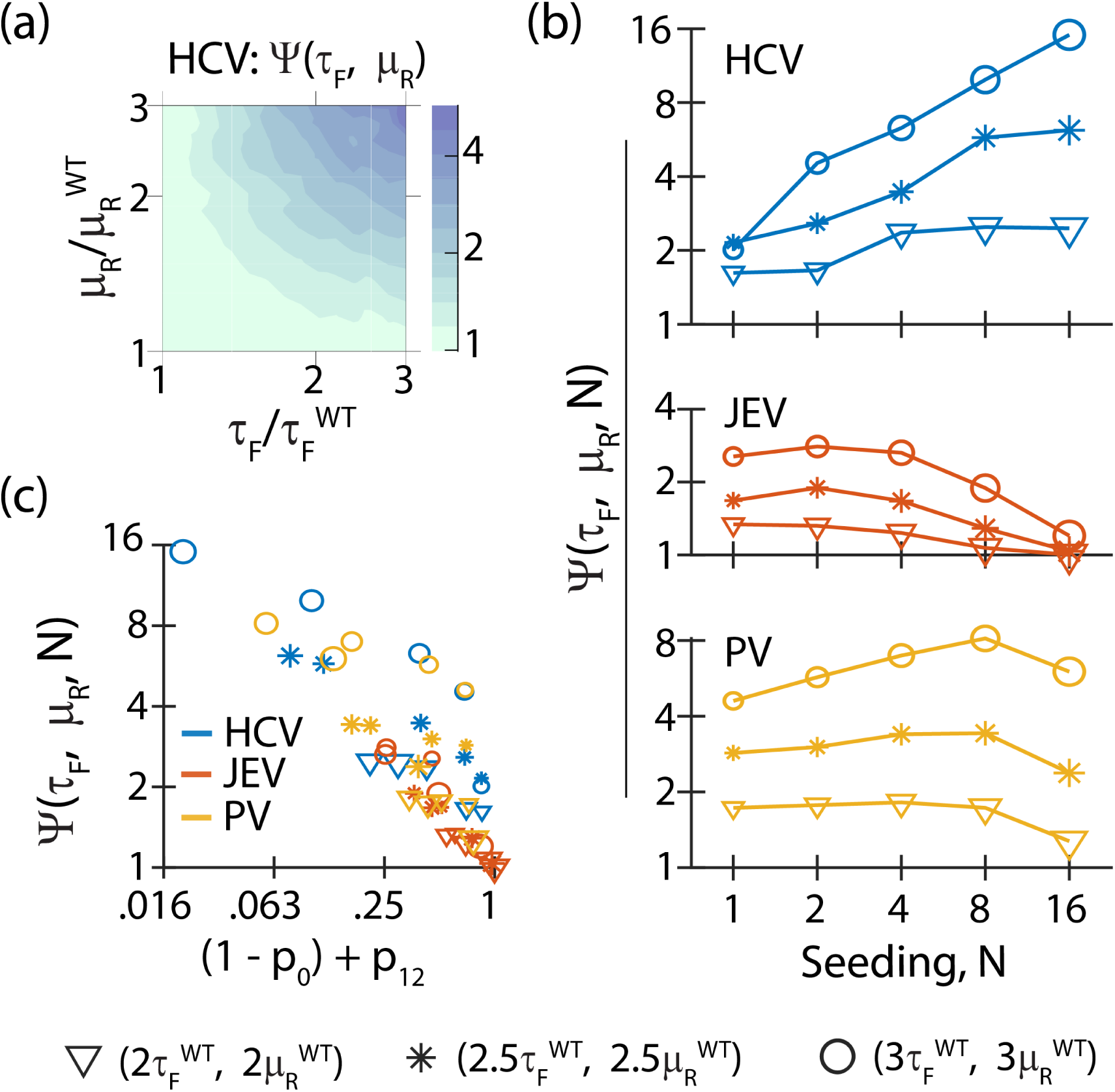
Synergy between life cycle processes affecting infectivity. (a) Ψ(*τ*_*F*_, *µ*_*R*_) evaluates the Bliss synergy between *τ*_*F*_ and *µ*_*R*_ for Φ, for HCV at seeding, N = 3. (b) Variation of Ψ(*τ*_*F*_, *µ*_*R*_) for various fold change in parameter values (denoted by different markers) with viral seeding (N) for each virus is shown. (c) Ψ(*τ*_*F*_, *µ*_*R*_) shows a negative correlation with {(1−*p*_0_)+*p*_12_}. 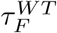 and 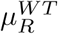 corresponds to the estimate for the respective virus. Marker properties are same in (b) and (c) and marker size corresponds to value of N.

At first glance, the quantitative relationship for *τ*_*F*_ − *µ*_*R*_ synergy varies with the virus-host system and seeding density in a complicated fashion (Figs. 5b). Under conditions where Φ → 1 or is close to extinction of infection (Φ → 0), perturbations do not influence Φ, individually or in combination. However, figure 5c shows that 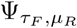 decreases with {(1 − *p*_0_) + *p*_12_}, a surrogate for how far the system is from either of the two deterministic limits (SI S4), where *p*_0_ and *p*_12_ denote Φ in unperturbed and doubly perturbed conditions, respectively. Similar synergy and associated negative correlation with {(1 − *p*_0_) + *p*_12_} is also predicted for *τ*_*F*_ − *k*_*t*_ (Figure S12c-g). Therefore, interventions that target membrane reorganization can be combined with other antiviral inhibitors to target early life cycle events and achieve efficient viral clearance.

## Discussion

We incorporated the dynamics of replication compartment (CM) formation accompanying cellular infection into a simplified intracellular life cycle description for monopartite (+)RNA viruses. This allowed us to capture observed dynamics for viruses spanning multiple (+)RNA virus families, parse diverse effects of host cell susceptibility, host factor silencing and virus mutations, as well as identify stochasticity associated with establishment of (+)RNA virus infection upon cell entry.

Based on temporal dependencies among model parameters, the viral life cycle can be categorized into three phases - establishment, growth and saturation (steady state). High translation efficiency (5′cap-dependent ribosome loading) and fast CM formation (associated with faster protein production), as observed for Japanese Encephalitis virus (JEV), results in rapid completion of the establishment phase. Following the compartmentalization of replication, the life cycle enters the growth phase that is marked by positive feedback from the newly synthesized RNA fueling the replication process. Kinetics of replication (*k*_*r*_), (+)RNA export from CM (*k*_*e*_) and (+)RNA degradation in cytoplasm shape this phase, which is particularly short for PV, owing to its rapid replication and export. Virus generation (*k*_*a*_) for PV is also very fast compared to the *Flaviviridae* members, which we attribute to differences in virus structural complexity, assembly and egress mechanisms [70, 71]. Replicative fitness (partly defined by *k*_*r*_ and *N*_*C*_) and viral RNA stability 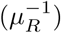 determines the steady state levels of cytoplasmic (+)RNA and viral titre in the final saturation phase for all viruses. Not surprisingly, *k*_*r*_, which is targeted via nucleoside inhibitors, remains a promising pan-viral drug target. Additionally, temporal sensitivity profiles suggests that the translation of PV is more sensitive to inhibition compared to replication while the reverse is true for JEV. Virus production is robust against reduction in assembly rate (*k*_*a*_) for all the three viruses considered here. Combined with steady state analysis, this suggests that genomes are packaged efficiently compared to their effective synthesis and degradation.

In our formalism, we also identify dynamics of CM formation (broadly defined by *τ*_*F*_ and *k*_*c*_) to be a key kinetic barrier in the early stage of the (+)RNA virus life cycle and it has been aptly described as the ‘load and choke point’ [17]. We demonstrate that ability of viruses to successfully establish infection in the host cell is stochastic and this ‘cellular infectivity’ (Φ) is determined at the onset of the infection. Such early stochastic extinction of viral infection has been similarly suggested due to biological noise [14, 13, 26]. Using our model based characterization of the viral life cycle, we are able to estimate this effect for the viruses. For synchronous co-infection, we predict that multiple genome infections are more likely to result in productive infection than single copy infection (as observed for PV [72]).

In a cell population, Φ quantifies the fraction of cells successfully infected upon entry of infectious viral particle(s) and correlates with the multiplicity of infection (MOI). As with MOI (discussed in [71]), we see that Φ also depends on seeding density (N) and virus-host determinants. Factors like viral genome stability in cytoplasm 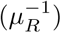 and kinetics of viral processes antecedent to formation of *RC*_*CM*_ affects productive infection. Indeed, drop in viral titers measured for translation-defective PV strains have been largely attributed to reduction in infectivity rather than growth defects [73]. Infectivity is highly sensitive to *τ*_*F*_ (compared to *k*_*t*_ or *k*_*c*_). This agrees with the reduction in Φ observed when membrane re-organization is hindered independently [6, 11]. Similarly, we speculate that the infectivity of a virus in a host cell can define its permissiveness [17, 74, 75].

Some of the early-infection parameters can also control cellular infectivity effectively in combination, displaying higher order effects due to their mutual action on common viral entities or processes. Our predictions are consistent with increased antiviral activity observed for membrane re-organizing inhibitor at lower MOI [12] as well as synergy observed between cell entry inhibitors and several classes of antiviral agents against HCV [67, 68]. Thus, strategies or drugs delaying CM formation, slowing translation, increasing viral degradation and reducing viral seeding, are expected to synergize in reducing infectivity. Host and viral heterogeneity would further accentuate this all-or-none dimorphism [15, 13].

Decrease in overall viral production due to lower infectivity can reduce viral seeding for the subsequent round of infections. Due to its dependency on viral seeding, such reduction in infectivity can manifest in a compounding effect that reduces the effective basic reproduction number, *R*_0_ and can lead to viral clearance. Therefore, cellular antiviral strategies that target cellular infectivity can be used in conjunction with other interventions (including action of innate immune response) that reduce the virus load.

Overall, our general theoretical framework can serve as a starting point for analysis of novel viruses with limited molecular level characterization, to generate insights into life cycle traits and bottlenecks, motivate design of experimental studies for insightful investigation and evaluate antiviral strategies.

## Methods

### Experimental data and model fitting

All experimental data sets used were curated from literature. Data for estimation of *τ*_*S*_ was retrieved from [37, 38, 39, 40]. Cellular life cycle dynamics data for HCV, JEV and PV were obtained from [22], [32] and [19], respectively. JFH1 (sgHCV strain) transfection dynamics and polyprotein dynamics of HCV NS4B mutants was obtained from [17] and [5], respectively. Effect of RTN silencing on viral dynamics were curated from [5, 7]. Figure digitization and data extraction were done using WebPlotDigitizer [76].

Estimation of parameter values was done using Iterative Approximate Bayesian computation (iABC) algorithm (SI SM1), which iteratively improves upon the distribution of parameter values based on *χ*^2^ statistics computed between model prediction and observed data, 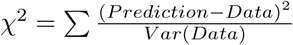 [77, 19]. In case variance was missing, we assumed a 25% relative error in the reported data and variance used in each case is shown in the figures. Practical identifiability is defined as the pairwise correlation in values of parameter combinations derived from the final iteration of estimation [19]. Further details are provided in the SI Methods.

### Temporal sensitivity analysis and estimation of Φ and Ψ

We used extended Fourier Amplitude Sensitivity Test (eFAST) [60, 78], to estimate the corresponding temporal sensitivity profile. Sensitivity indices for *RC*_*CM*_, *V*_*T*_ and *R*_*cyt*_, corresponding to a change of up to 10% in parameter values, were evaluated every 1.5 minutes through the course of the infection to generate the temporal profile.

To estimate cellular infectivity (Φ), stochastic realizations of the life cycle were implemented using the Gillespie algorithm [79], and classified as (a) successful infection (*I*_*S*_) if *RC*_*CM*_ is formed, (b) failed infection (*I*_*F*_) if all viral (+)RNA degrade, and (c) inconclusive if neither happen within a stipulated time. Since the fate of infection was decided in all stochastic realizations for HCV life cycle (slowest of the three viruses) by 12 h, this was used for the simulation run time. We consider only the ‘conclusive’ realizations and define Φ as 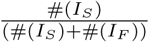. Dynamics of *f*_*CM*_ is incorporated by updating it at every event or at steps of 0.05*τ*_*F*_ (whichever is shorter). This limits the error due to discretization of *f*_*CM*_ to 6.5%.

To calculate synergy (Ψ), between two parameters, *a*_1_ and *a*_2_, that reduce Φ, we estimate *p*_0_, *p*_1_, *p*_2_ and *p*_12_ as Φ corresponding to no change, change in parameter *a*_1_, parameter *a*_2_, and in both parameters, respectively. Synergy among parameters is given by, 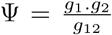 where *g*_*X*_ denotes 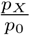 for *X* ∈ 1, 2, 12 (Bliss criterion [69]).

## Availability of code

Codes can be downloaded from our Github repository, https://github.com/hcharsh/iABC_fit/tree/master/%2BRNA_viral_lifecycle_fit.

## Acknowledgement

This work was supported by the Indian Institute of Science Bangalore (RR), Wellcome Trust—DBT India Alliance intermediate fellowship (RR), Council of Scientific and Industrial Research fellowship (VAR) and the Prime Minister Research Fellowship (HC). We thank Narendra Dixit, Mohit Jolly, Sunaina Banerjee and Suraj Jagtap for valuable feedback on the manuscript.

## Competing interests

The authors declare that they have no competing interests.

### Cellular life cycle model

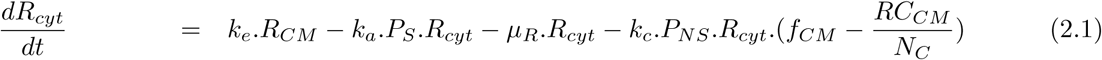

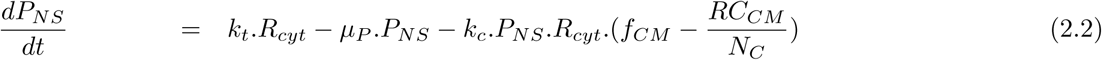

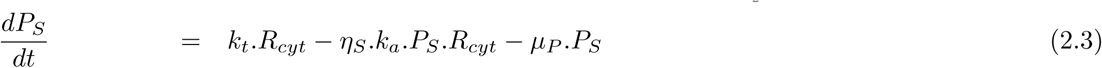

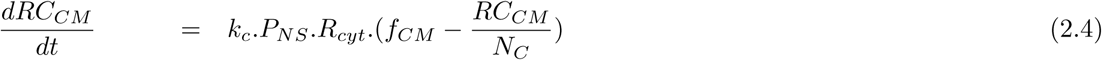

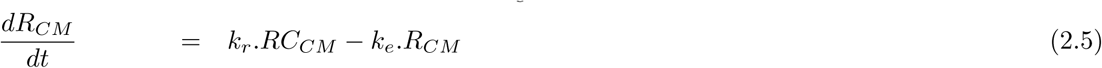

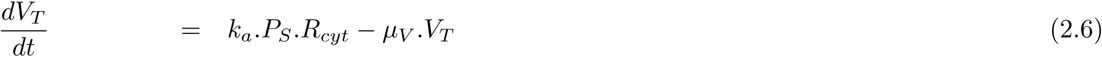

## Outline of the Supplementary Information file

**SI S1 Description of the model**

**SI S2 Revisiting the assumptions**

**SI S3 Steady state analysis**

**SI S4 Quantifying “far from determinstic regime”**

**SI S5 Alternate models**

**SI SM1 Iterative Approximate Bayesian computation (iABC) for parameter estimation**

**SI SM2 Practical Identifiability calculation**

**SI SM3 Implementation details**

**Figure S1 Compartment formation dynamics**

**Figure S2 Parsing effects of host cell permissiveness**

**Figure S3 Fitting transfection dynamics using different fitting options**

**Figure S4 Algorithm convergence and practical identifiability analysis for fitting cellular dynamics of HCV, JEV and PV using** 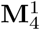

**Figure S5 Analysing alternate model, M**′: *f*_*CM*_ **formulation variant where** *f*_*CM*_ = 1

**Figure S6 Analysing alternate model**, 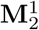: *f*_*CM*_ **formulation variant where** 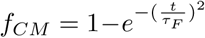

**Figure S7 Analysing alternate model**, 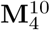**: formulation assumes that 10** *P*_*NS*_ **molecules are required to form one** *RC*_*CM*_

**Figure S8 Analysing alternate model, W**_4_**: Packaging formulation variant where genomes from compartments associate with SP from cytoplasm to form new viral particles**

**Figure S9 Parameter-temporal sensitivity profiles for dynamics of (+)vRNA in cytoplasm (***R*_*cyt*_)

**Figure S10 Cellular infectivity profiles**

**Figure S11 Seeding dependent fold change in** Φ **due to change in life cycle parameter value**

**Figure S12 Synergestic strategies to reduce cellular infectivity**

**Table S1 Compartment formation dynamics: Parameter estimation and goodness of fit**

**Table S2 Cellular life cycle model: Parameter estimation summary**

**Table S3 Comparing parameter estimation for sgHCV (JFH1 strain) life cycle in host cells of different permissivity**

**Table S4 Comparing parameter estimation and goodness of fit for model variants**

## Supplementary Information for

### SI S1 Description of the model

To capture the viral dynamics with a generalized model, we focus on life cycle processes common to all monopartite +ssRNA viruses. Virus enters the cell and uncoats to release its genome, +RNA, into the cytoplasm. This is usually a fast process relative to the corresponding viral cellular infection span, as suggested by direct kinetics measurement [5] or detection of early protein production (a post ‘un-coating’ event) [1]. Thus for a reduced model, we do not explicitly account for this fast step of viral entry. Furthermore, we coarse-grain over specific molecular details to keep model formulation simple. For example, we ignore the specific molecular roles of individual viral proteins. We use a general term for non-structural proteins (*P*_*NS*_) that are involved in the replication process and one for structural proteins (*P*_*S*_) which are required for viral assembly.

#### Viral protein synthesis

In the cytoplasm, the viral +RNA strands associate with host ribosomes, to undergo translation. Across the class of the +RNA viruses, different ribosome loading mechanisms has been observed for viruses [25, 31, 21], which can influence the kinetics and the number of ribosomes simultaneously loaded on a translating RNA strand. Translation produces polyprotein that is co-and post-translationally processed (by host and viral proteases) to produce individual viral proteins (*P*_*NS*_ and *P*_*S*_). This entire process of protein synthesis is simplified by lumping it from ribosome loading to polyprotein cleavage into a single kinetic step.

Assuming abundance of host ribosome, the process is modelled as a first order reaction proportional to level of +RNA strands in the cytoplasm, *R*_*cyt*_ (eqs. 2.2 and 2.3). The rate parameter, *k*_*t*_, accounts for rate of single event of protein production from *R*_*cyt*_ multiplied by the average number of such simultaneous events occurring on each cytoplasmic viral +RNA strand.

Previous computational studies [9, 1] suggest that ribosome loading and translation elongation steps are fast compared to polyprotein cleavage for HCV, thus it is likely that polyprotein processing is the rate determining step and defines the parameter *k*_*t*_.

#### Compartment formation

Cellular life cycle of +RNA viruses are marked by spatial segregation of viral replication from the host cytoplasm. During infection, viral non-structural proteins induce morphological changes in the membranes of host organelle (target organelle varies with virus) to form of replication compartments (CMs) [24, 26, 10, 19, 34]. These compartments confine (viral) replication substrates and protect them from cytosolic degradation.

Although a critical life cycle event for +RNA viruses and probably a rate limiting process, formation dynamics of CMs has not been given due attention in previous models [9, 4, 1, 37]. To account for the kinetics of CM development, we use a data driven approach augmented by a reasonable assumption. Though compartment formation is induced by *P*_*NS*_ and we don’t have sufficient quantitative data to reliably account for the correlated dynamics. Therefore, we assume that CM formation is triggered by viral entry. Furthermore, it is difficult to independently measure the functional development of CM that imparts protective confinement of replication. So, we estimate the rate parameter via life cycle fitting. However, the mathematical form of the dynamics is assumed to be identical to structural development of CMs, which has been quantitatively monitored for various +RNA viruses. Our analysis show that a Weibull function with exponent of 4, captures the structural manifestation quite accurately (Figure 1, main text; Table S1). Thus we assume the same form, given by 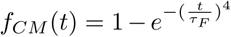 (eq. 1, main text), to model the functional maturity of CMs. In the expression, *τ*_*F*_ is the delay parameter associated with the dynamics.

An infection-triggered induction, rather than a *P*_*NS*_ triggered induction, reduces the number of free parameters. Furthermore, it allows us to implicitly lump time lags associated with entry and un-coating, into *τ*_*F*_. However these sub-events can be explicitly modelled in future with more measurements.

#### Compartmentalization of replication intermediate

Viral *R*_*cyt*_ associates with *P*_*NS*_ and unoccupied compartments to form compartmentalized replication complexes (*RC*_*CM*_). Given the limited number of unoccupied compartments available at a given time, the compartmentalization process can be viewed as a competition among complexes to be encapsulated. The dynamics of *RC*_*CM*_ formation is modelled as a logistic function with the growth rate proportional to *R*_*cyt*_ and *P*_*NS*_ and carrying capacity being proportional to level of unoccupied compartments at that time, given by 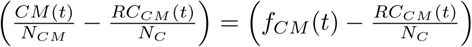.

The rate constant, *k*_*c*_, captures the compartmentalization kinetics whereas *N*_*C*_ denotes the number of compartmentalized replication complexes sustained during the infection (eqs. 2.1, 2.2 and 2.4, in Main text). Both *N*_*C*_ and *k*_*c*_ depend on stability of the replication complex in the host cell. Additionally, *N*_*C*_ is also influenced by the number of replication compartments available, and *k*_*c*_ is influenced by factors like cis-replicating element, compartment accessibility and genome RNA-*P*_*NS*_ association kinetics. Defining *N*_*CM*_ as the carrying capacity of replication compartments formed during the course of infection, we can say that on an average 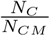 replication intermediates occupy a given compartment.

#### Compartmentalized replication and +RNA export from compartments

Inside a compartment, new viral +RNA strands are synthesized at a rate proportional to number of replication complexes compartmentalized into the compartment. We assume most +RNA synthesis occurs inside the compartments, ignoring cytoplasmic production of +RNA in comparison. Hence, the total rate sould be proportional to total *RC*_*CM*_ present in the cell at that time (eq. 2.5). The rate constant, *k*_*r*_, denotes the net +RNA synthesis rate per *RC*_*CM*_, determined by replication elongation kinetics as well as the (average) number of simultaneous replication events occurring on each *RC*_*CM*_.

In the model, we assume a temporally averaged value for *k*_*r*_ which is a good approximation. The number of viral replicase is fairly constant as they are protected from degradation inside the compartments [24, 26, 10] and the import of additional viral replicase protein into a compartment drops drastically with the age of the compartment [35]. Furthermore the viral +RNA present in the compartment are exported into the cytoplasm. The process is modelled as a first order reaction (eqs. 2.1, 2.5). The rate is proportional to +RNA in compartments (*R*_*CM*_) and parameterized by *k*_*e*_.

#### Virus generation

Viral genome associate with various structural proteins *P*_*S*_ to form a viral particle, which exits the cell either via a lytic or lysogenic mode, depending on the virus. Recent studies show that assembly occurs close to the compartmental structure, and is phase separated from the cytoplasm [7]. In addition to the viral egress mechanism, the mechanism of viral assembly also differs with virus [30]. Though a lot has been discovered, current understanding of precise mechanism is far from complete. This, coupled with our attempt to simplify the life cycle model, we coarse-grain the description of the entire viral generation process with a single equation.

Virus generation process is modelled as a second order reaction between *R*_*cyt*_ and *P*_*S*_ with rate constant, *k*_*a*_, to produce virus particles (*V*_*T*_) (eqs. 2.1, 2.3 and 2.6, Main text). The parameter, *k*_*a*_, is affected by assembly mechanism, genome-capsid association kinetics as well as viral egress mode. Further, cis-acting elements and packaging signal sequence on the genome can also modulate *k*_*a*_. Additionally, *η*_*S*_ accounts for stoichiometry of viral particle composition. *η*_*S*_ copies of each structural protein associate with one genome to form one virus particle.

#### Degradation

Viral genome and proteins are susceptible to degradation in cytoplasm (eqs. 2.1, 2.2, 2.3). Motivated by experimental data[1], we also consider degradation of virus (eq. 2.6). However, we ignore the degradation of viral molecules inside compartment, considering the protective environment inside the CMs [24, 26, 10]

### SI S2 Revisiting the assumptions

We have tried to keep the model simple, so as to get reliable estimates for model parameters with limited experimental data available. Furthermore, lumping mechanistic and molecular details, makes the model applicable to a wider range of viruses. Such assumptions are discussed here, which can be challenged in future with better understanding and/or more quantitative data.

In our formulation, we assume that CM formation is triggered by the onset of infection, not by protein level (*P*_*NS*_). One could incorporate the effect of *P*_*NS*_ and model CM formation with an activating Hill function. For example, 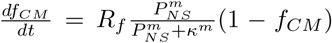, where *κ* and m are the threshold and shape parameters of the Hill function, respectively. *R*_*f*_ represents the maximal rate of CM formation, when *P*_*NS*_ *>> κ* and *f*_*CM*_ *<<* 1. However in this formulation, the number of free parameters increases, which without independent estimation(s) will lead to over-fitting of data. Thus given the limited data, we chose the Weibull function with a one time and one steepness parameter.

In our model, viral entry and un-coating delays are implicitly lumped with the true CM formation lag within *τ*_*F*_. However, one should note that it is possible that viral +RNA degradation rate may vary depending on its state of virus encapsidation. Thus, subsequent models can consider such sub-events more carefully. A small (but non-zero) basal level of +RNA synthesis occurs in the cytoplasm outside the CM. However, the contribution is small compared to replication occurring in the CM. In case of HCV, it is known that mutants defective in inducing CM formation [36] have dynamics similar to replication deficient mutants[4], Although we have not included it explicitly, the effect of cytoplasmic replication is lumped in the degradation rate of viral +RNA in cytoplasm (*µ*_*R*_). We have also ignored low level of degradation of viral +RNA and replication complexes inside CMs, based on experimental evidence.

Inside the compartments, we do not explicitly account for formation of dsRNA due to replication of nascent +RNA inside the compartments, rather +RNA have to be exported before they can synthesize a replication intermediate. This simplifying assumption works well in our deterministic mean field setting owing to the high bias of replicase enzyme to replicate the -RNA strand over the +RNA strand [2, 9].

The formation of compartmentalized replication complex (*RC*_*CM*_) has been depicted as a single step association between *R*_*cyt*_, *P*_*NS*_ and unoccupied compartments. The assumption is implicit in the temporal order of the events happening, that is, it does not differentiate whether (a) *R*_*cyt*_ and *P*_*NS*_ associate first and then enter the unoccupied RC, or (b) *R*_*cyt*_ and *P*_*NS*_ enter the unoccupied RC and then associate. Furthermore the process of association of NS proteins with +RNA to form replication complexes can be divided into multiple steps, of which some are cis-mediated (like cis-replication element activity, NS3-5B protein association) whereas others are trans-mediated [14, 18]. Cis-mediated reactions can be modelled as pseudo-first order reactions, and the trans-mechanisms by second order reactions. Further assuming that a trans-mediated sub-process is the rate limiting step in complex formation [9, 4, 1, 37], and that the complex also associates with the unoccupied compartments, we model the kinetics of +RNA-*P*_*NS*_-’unoccupied CM’ association to form compartmentalized replication complex as a third order reaction. The rate kinetics of the other fast sub-processes are lumped in the kinetic parameter, *k*_*c*_.

Similarly the process of viral particle generation constitutes of multiple sub-processes, kinetics of which have been lumped into *k*_*a*_ (see previous section, SI S1). Future models could consider this process with higher granularity.

### SI S3 Steady state analysis

Steady state conditions for non-zero viral seeding:

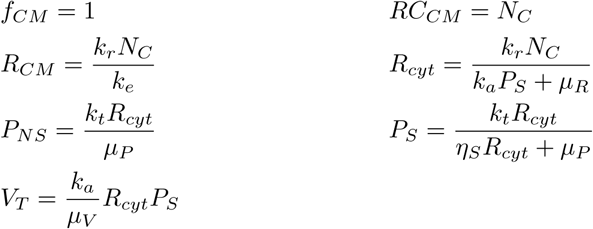

Simplification under various limits:

1. When *k*_*a*_*P*_*S*_ *>> µ*_*R*_, *µ*_*P*_ we have 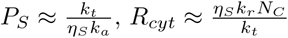 and 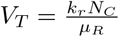.
2. When *k*_*a*_*P*_*S*_ *<< µ*_*R*_, *µ*_*P*_ we have 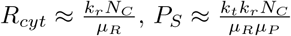 and 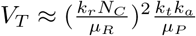.
3. When the two processes (assembly and cytoplasmic degradation of +RNA or *P*_*S*_) are comparable then *V*_*T*_ is super-linear but sub-quadratic with increase in +RNA levels. In this regime, an increase in *k*_*a*_*P*_*S*_ due to higher *k*_*t*_ leads to comparatively lower increase in *R*_*cyt*_ (and thus *P*_*NS*_) due to increase in *N*_*C*_. This explains the trends between *P*_*NS*_ and +RNA observed for HCV and JEV.

### SI S4 Quantifying “far from deterministic regime”

We define *p*_0_ and *p*_12_ as the cellular infectivity corresponding to no perturbation and combined perturbation respectively. Let us consider individual perturbations which reduce Φ, so *p*_0_ ≥ *p*_12_. Then, the following situations are possible

a. *p*_0_ = 0 ⇒ *p*_12_ = 0. In either situation, unperturbed or perturbed, the infection will go extinct - the deterministic infection extinction limit.
b. *p*_12_ = 1 ⇒ *p*_0_ = 1, implying that viral seeding always results in a productive infection - the deterministic infection sustenance limit. Here these limits are defined for a particular system, characterized by the virus-host pair, seeding density and level of parameter change, being considered.

So we see that (1 − *p*_0_) and *p*_12_ positively correlate with how far system is from the limit of deterministic infection extinction and sustenance, respectively. Hence, we use {(1 − *p*_0_) + *p*_12_} as a first-order approximation to quantify how “far the system is from either of the deterministic regimes”.

### SI S5 Alternate models

#### Formulations

An important difference between the our proposed model and previous models existing in literature is that we incorporate the dynamics of compartment formation. We assume that the normalized dynamics of functionally active compartment increases as 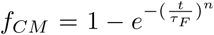, (eq. 1) with n = 4 as obtained from the considering the structural development of compartments; this model is referred to as 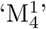. Here, we explore two alternative formulations of *f*_*CM*_.

1. pre-formed CM present: *f*_*CM*_ = 1 (equivalent to earlier models [9, 1]). This model is referred to as M′.
2. *f*_*CM*_ is described by Weibull function but with exponent, n = 2. Hence for this alternate model, 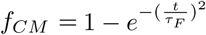. This model is referred to as 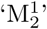.

Note as *τ*_*F*_ → 0, 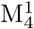 (or 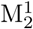) converges to M′; thus M′ can be considered as a special case of 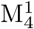 (or 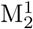).

Another alternate formulation stems from the stoichiometric consideration for the replication complex. In our model we assume that one *P*_*NS*_ and one *R*_*cyt*_ are used to form one *RC*_*CM*_. Thus, we explore a scenario (third alternate model) where ten *P*_*NS*_ and one *R*_*cyt*_ are used to from one *RC*_*CM*_. This alternate model is referred to as 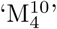.

For this model, the only equation different from 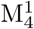 is eq. 2.2, which is modified to 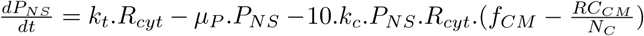.

However, since only a tiny fraction of NSP formed localize in the compartments, we did not expect the stoichiometry to significantly affect the life cycle dynamics.

Furthermore, in 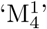, viral production uses genome from the cytoplasm since structural proteins are also present there. However some reports indicate that assembly is also spatially segregated from the cytoplasm, and may be linked with replication compartments [26, 10, 11, 7]. Thus we consider an alternate model (the fourth alternate) for virus generation, where genome from compartment (*R*_*CM*_) associates with *P*_*S*_ to form virus; this is similar to the model presented by Aunins *et al*, 2018 [1]. This is referred to as W_4_.

However this model (W_4_) and the one presented by Aunins *et. al* [1], do not explicitly locate where capsid genome assembly occurs. This formulation suggests that the structural proteins enter into the compartments where genome were formed, and the import kinetics is lumped into the assembly rate. Nevertheless, this indicates a directionality among the viral processes, which one may argue not to be the optimal strategy for the virus [23]

#### Compartmental viral +RNA packaging model (W_4_)

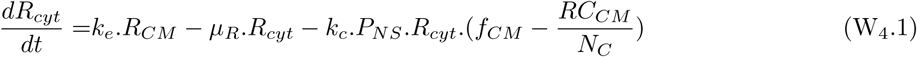

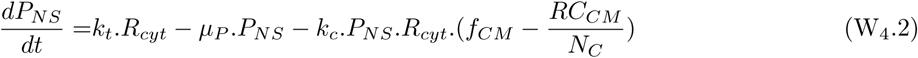

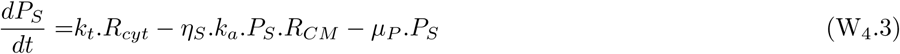

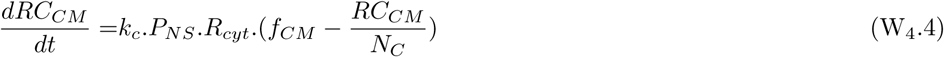

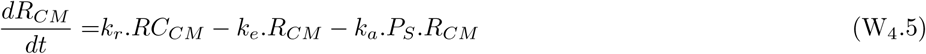

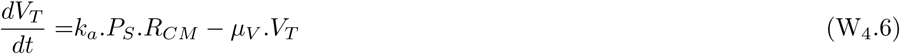

#### Comparison

In addition to the main model 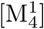, we fit the four alternate models to the observed cellular dynamics of viral life cycle of HCV in Huh7 cells [1], JEV in PS cells [32] and PV in HeLaS3 cells [28] and estimate the values of model parameter (Figs S5, S6, S7 and S8). Fitting statistics are summarized in Table S4. Based on AIC values for each model, cumulated over analyses of three infection systems, 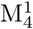 and M′ seem to be slightly better than others (*δAIC* ≈ 2). Similar AIC values for 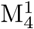 and M′ inspite of one extra parameter in 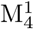 demonstrates the merit of our model.

We see that accounting for CM formation (as in 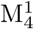) better fits the dynamics of viral life cycle when CM formation delay and infection span are comparable (as in PV). 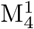 estimates about 88 HCV compartmentalized replication complexes (*N*_*C*_, Table S2), which matches more closely with experimentally reported range of 70-100 [15, 4, 13], compared to the corresponding estimate by M (*N*_*C*_ ≈300, Table S4). Further the steady state ratio of +RNA to -RNA predicted by 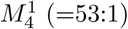, about 1.6 times the experimental observations [35, 12] whereas the (corresponding) ratio predicted by M′ (=12.5:1), more than 2-fold smaller (than the observations). Thus, 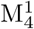 can be considered a better generalization compared to M′, to model cellular life cycle of +ssRNA viruses.

### SI SM1 Iterative Approximate Bayesian computation (iABC) for parameter estimation

We implement an iterative version of Approximate Bayesian computation (ABC)[3, 28] for data fitting. We iteratively improve upon distribution of each parameter, based on *χ*^2^ statistics of parameter sets sampled using them. The algorithm used is as follows:

1. Step 1: Initial guess for parameter estimate distribution
2. Step 2: *V* parameter sets are sampled using the current estimate of the distribution. Calculate *χ*^2^ statistics for each of the parameter sets.
3. Step 3: Sort the parameter sets based on their *χ*^2^ statistics. Select *M* parameter sets with lowest *χ*^2^ statistics.
4. Step 4: Use the *M* parameter sets to calculate the new estimate for the distribution. Repeat Steps 2 through 4 until stopping condition is satisfied.

Latin hyper-cube sampling (LHS) [22] is used to sample parameter sets from the distribution. To improve robustness of the algorithm and not spiral into local minima, we sample parameter combinations (in Step 2) from current estimate of distribution combined with a uniformly distributed noise.

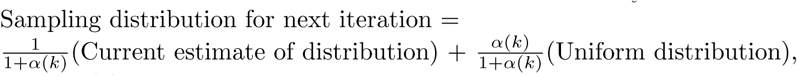

where *α*(*k*) is the strength of perturbation, which decreases with the index of iteration, *k*.

Stopping conditions can be based on numerical convergence in successive estimates of parameter distribution or on *χ*^2^ statistics. In our implementation, the algorithm was stopped after a fixed number of iterations, *k. k* was chosen such that the difference between the largest *χ*^2^ statistics of the parameter sets used for estimating the new distribution converge (Fig S4 a, b, c).

### SI SM2 Practical Identifiability calculation

We calculate the practical identifiability of parameter estimation to characterize the redundancy in parameter distribution estimated using the given data. This is done by evaluating the correlation among the values of different parameters which form the parameter sets used to calculate the distribution [28].

Suppose 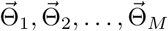, are the M parameter sets selected based on *χ*^2^ statistics. We calculate the correlation between 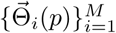 and 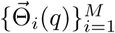 to characterize the redundancy between the estimates of the *p*^*th*^ and *q*^*th*^ parameters.

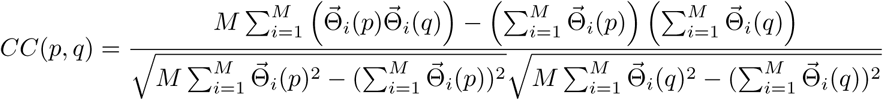

Note that *CC*(*p, q*) is defined when *p* ≠ *q* and the correlation matrix, as expected, is symmetric, that is, *CC*(*p, q*) = *CC*(*q, p*).

Furthermore, lower the absolute value of *CC*(*p, q*) - correlation coefficient between the *p*^*th*^ and the *q*^*th*^ parameters - higher is the pair-wise identifiability between the estimates of the parameter given the data set.

### SI SM3 Implementation details

The ODE models were solved using MATLAB 2019b ODE solver, ‘ode23s’. Codes for the analysis can be downloaded from our Github repository, https://github.com/hcharsh/iABC_fit/tree/master/%2BRNA_viral_lifecycle_fit. In addition to the data, we need to specify the fixed parameters and initial conditions (if known) to define the fitting problem.

We fix parameter values associated with various degradation rates and viral particle stoichiometry either based on literature survey or independent data analysis, indicated in this section. We also list the initial guesses (for free parameters), algorithm options and initial conditions (if any) used in the study. In our analysis we fit the logarithm of parameter values (normalized to the units mentioned in Table 1, Main text) in base 10, to explore a large dynamic range for each parameter value.

In this section, *u*[*a, b*] denotes the uniform distribution with lower and upper bounds as *a* and *b* respectively.

### SI SM3.1 Characterizing ultra-structural data corresponding to formation of replication compartments

**Initial guess** for the parameter distribution for fitting compartment formation dynamics (eq. S1) observed in all four infection systems:

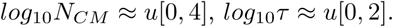

#### iABC algorithm options

k = 5; V = 1 × 10^4^; M = 0.025N; *α* = 0.25 for all of the systems considered here.

### SI SM3.2 Characterizing the intracellular infection dynamics of HCV in Huh7, JEV in PS cell or PV in HeLaS3

#### Fixed parameters

*η*_*S*_ denotes the number of copies of each structural proteins required to form a viral particle. For HCV: *η*_*S*_ = 180[29]; For JEV: *η*_*S*_ is assumed to be same as that of DENV[17] or HCV, so *η*_*S*_ = 180; For PV: *η*_*S*_ = 60 [20]. For HCV, JEV and PV: cytoplasmic degradation rate of viral +RNA, *µ*_*R*_ = 0.25 h^−1^[1]

For HCV, JEV and PV: cytoplasmic degradation rate of viral proteins, *µ*_*P*_ = 0.11 h^−1^[1]

For HCV, JEV and PV: degradation rate of extracellular viruses, *µ*_*V*_ = 6 × 10^−2^ h^−1^ - estimated using extracellular virus dynamics observed (till 15 hours post infection) for HCV [1].

**Initial condition** used (all variables start at zero unless specified here):

For the model and all its variants, we have

For HCV: According to MOI specified [1], *R*_*cyt*_(0) = 3 and *P*_*S*_ = *η*_*S*_*R*_*cyt*_ = 540. Based on data [1], *V*_*T*_ (0) = 4.6.

For JEV: According to MOI specified [32], *R*_*cyt*_(0) = 10. Based on data [32], *V*_*T*_ (0) = 0.01.

For PV: According to MOI specified [28], *R*_*cyt*_(0) = 10.

**Initial guess** for the parameter distribution:

*log*_10_*k*_*t*_ ≈ *u*[0, 3], *log*_10_*k*_*c*_ ≈ *u*[−5, −1], *log*_10_*τ* ≈ *u*[0, 2], *log*_10_*k*_*r*_ ≈ *u*[−1, 3], *log*_10_*k*_*e*_ ≈ *u*[−2, 2], *log*_10_*N*_*C*_ ≈ *u*[1, 4], *log*_10_*k*_*a*_ ≈ *u*[−10, 0].

#### iABC algorithm options

k = 8; V = 1 × 10^4^; M = 0.025N; *α* = 0.25 for all three infection systems for all the model variants.

### SI SM3.3 Note on fixing the same value for cytoplasmic viral RNA degradation rate for HCV and JEV

Parameter estimation (Table S2) suggests that *τ*_*F*_ = 2.6*h* for JEV whereas it is 5.8*h* for HCV. Faster protein production (higher *k*_*t*_) leading to a rapid, early increase in NSP levels and consequently an early trigger for membrane re-organization most likely explains the smaller lag in RC formation for JEV. Even differences in re-organization mechanisms, across the different viruses, may contribute too.

However the observed trend may also be an artefact of our implementation procedure: here we assume *µ*_*R*_ = 0.25*h*^−1^ for all infection systems. Studies suggest that 5’capping increases stability of viral RNA[6]; and the low value of *τ*_*F*_ estimated may be compensating for the fixed parameter value used; as smaller *τ*_*F*_ or smaller *µ*_*R*_ are functionally similar.

### SI SM3.4 Accounting for Luciferase dynamics to analysis transfection dynamics

We use the following equation to account for the dynamics of Luciferase while estimating parameters for (a) sgHCV transfection in Huh7-Lp and Huh-Lunet cell lines [4], and (b) life cycle of WT and NS4B mutant HCV strains [36]:

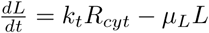

Here L denotes level of Luciferase, produced upon translation of *R*_*cyt*_ (viral +RNA in cytoplasm). *µ*_*L*_ denote the degradation rate of Luciferase which is fixed at 0.44*h*^−1^ based on experimental evidence reported [4].

### SI SM3.5 Characterizing the intracellular transfection dynamics of JFH1 (sgrHCV) in Huh7-Lp and Huh7-Lunet cell lines

#### Fixed parameters

*µ*_*R*_ = 0.25 h^−1^[1]

*µ*_*P*_ = 0.11 h^−1^[1].

*µ*_*L*_ = 0.44 h^−1^[4].

Further assembly and viral degradation terms are ignored as sgHCV strain can not form new viral particles, *k*_*a*_ = 0 and *µ*_*V*_ = 0.

**Initial condition** used: *R*_*cyt*_(0) = 100 and all other variables start at zero.

**Initial guess** for the parameter distribution:

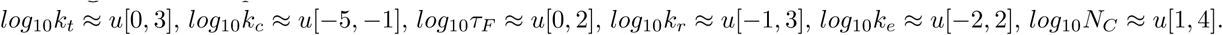

**iABC algorithm options**: k = 3; V = 5 × 10^3^; M = 0.025N; *α* = 0.25 for both the transfection systems.

The analysis is done assuming 25% constant relative error in data reported (Fig. S2), as the range of reported error varies a lot [4]. However we also fit the model using *R*_*cyt*_(0) = 100 using reported experimental error [4] and *R*_*cyt*_(0) = 1000 assuming 25% constant relative error. The estimation is summarized in Fig. S3.

### SI SM3.6 Intracellular life cycle of sgHCV NS4B mutants

*τ*_*F*_ is the only parameter being varied while fitting normalized protein dynamics observed for sgHCV NS4B mutant strain. We fix the value of the remaining parameters according to those estimated in Table S2 (HCV infection dynamics in Huh cells). *τ*_*F*_ is kept fixed for wildtype strain (Table S2). Since we do not know the initial condition (viral seeding) precisely, we fit for it using the normalized protein dynamics observed for wildtype strain. The viral seeding, thus obtained, is used for the analysis of *τ*_*F*_ for the mutant strains. Additionally we fix *µ*_*L*_ at 0.44 h^−1^ [4].

**Initial guess** for fitting WT dynamics: *log*_10_[*R*_*cyt*_(0)] ≈ *u*[0, 2],

*R*_*cyt*_(0) estimation: Median=12.45; first quantile=11.8; third quantile=13.6.

**Initial guess** for fitting mutant dynamics: *log*_10_*τ*_*F*_ ≈ *u*[0, *log*_10_144].

#### iABC algorithm options

k = 2; V = 10^3^; M = 0.025N; *α* = 0.25 for each fitting exercise.

### SI SM3.7 Note

While fitting JEV life cycle dynamics, the normalized dynamics of *RC*_*CM*_ was used to explain time series measurements of relative RdRp activity, and steady state of *RC*_*CM*_ was used to fit the steady state of (-)vRNA. This is because *RC*_*CM*_ characterizes active replication. However in case RdRp activity measurement is not available, we fit *RC*_*CM*_ dynamics to that of (-)vRNA. Furthermore we fit dynamics of (*R*_*cyt*_ + *R*_*CM*_ + *RC*_*CM*_) to that of (+)vRNA, as in our model +RNA is present in cytoplasm (*R*_*cyt*_) and in compartment either free (*R*_*CM*_) or as part of the dsRNA replication intermediate (*RC*_*CM*_). Total vRNA considers levels of both (-)vRNA and (+)vRNA strands.

**Figure S1:**
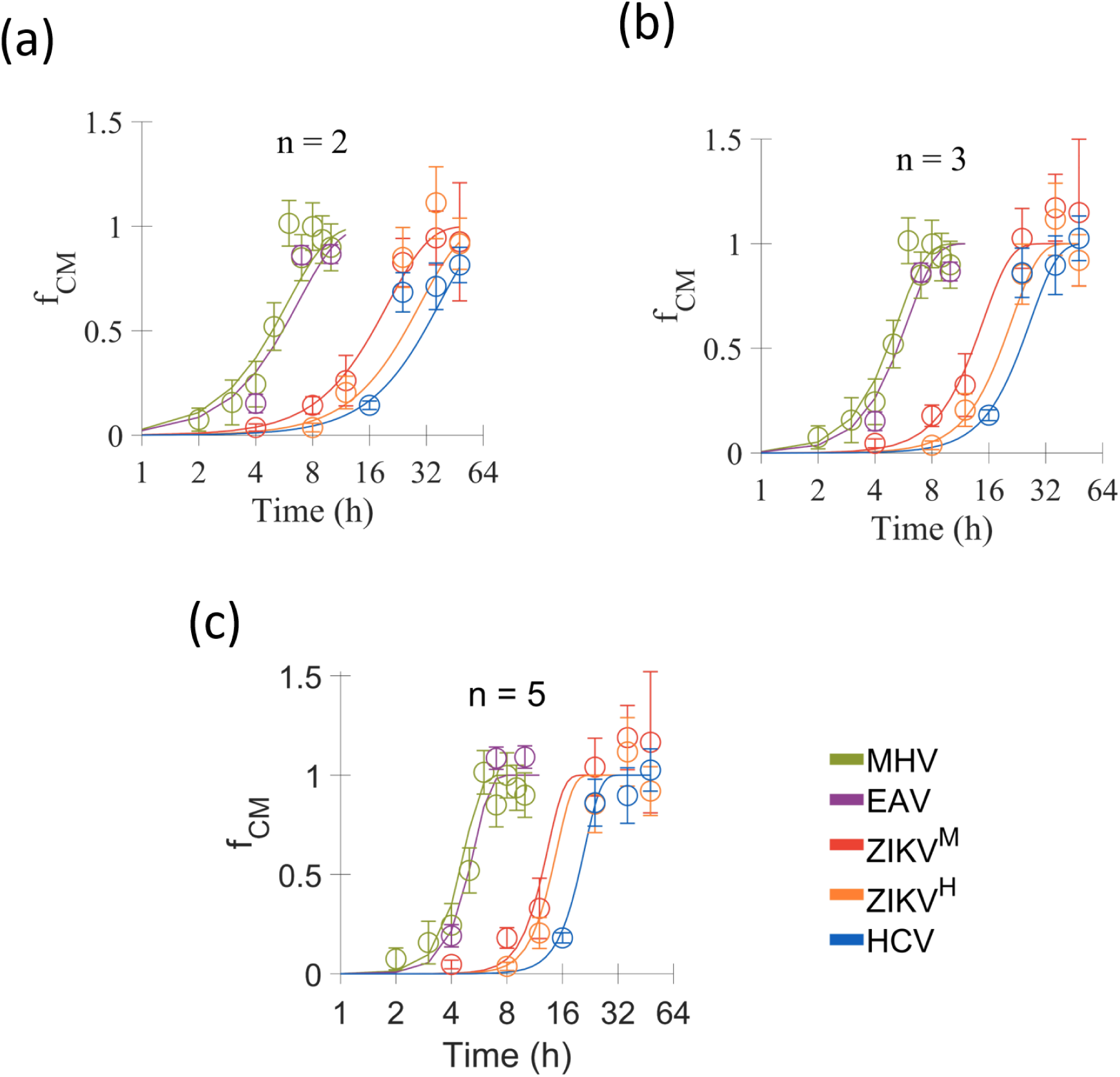
Compartment formation dynamics. Normalized dynamics of compartment formation (*f*_*CM*_) observed for different (+)RNA viruses and fit (lines) using eq. S1 with (a) n = 2, (b) n = 3 and (c) n = 5. ZIKV^*M*^ and ZIKV^*H*^ denotes the MR766 and H/PF/2013 strains of Zika virus, respectively. Data derived from [16, 33, 27, 8]. Thin (lightly colored) lines represents dynamics predicted using a set of best parameter combinations from iABC (see section SI SM3 for algorithm implementation details) and thick lines denotes their average. Here, hollow circles and error bars correspond to data used for fitting.

**Figure S2:**
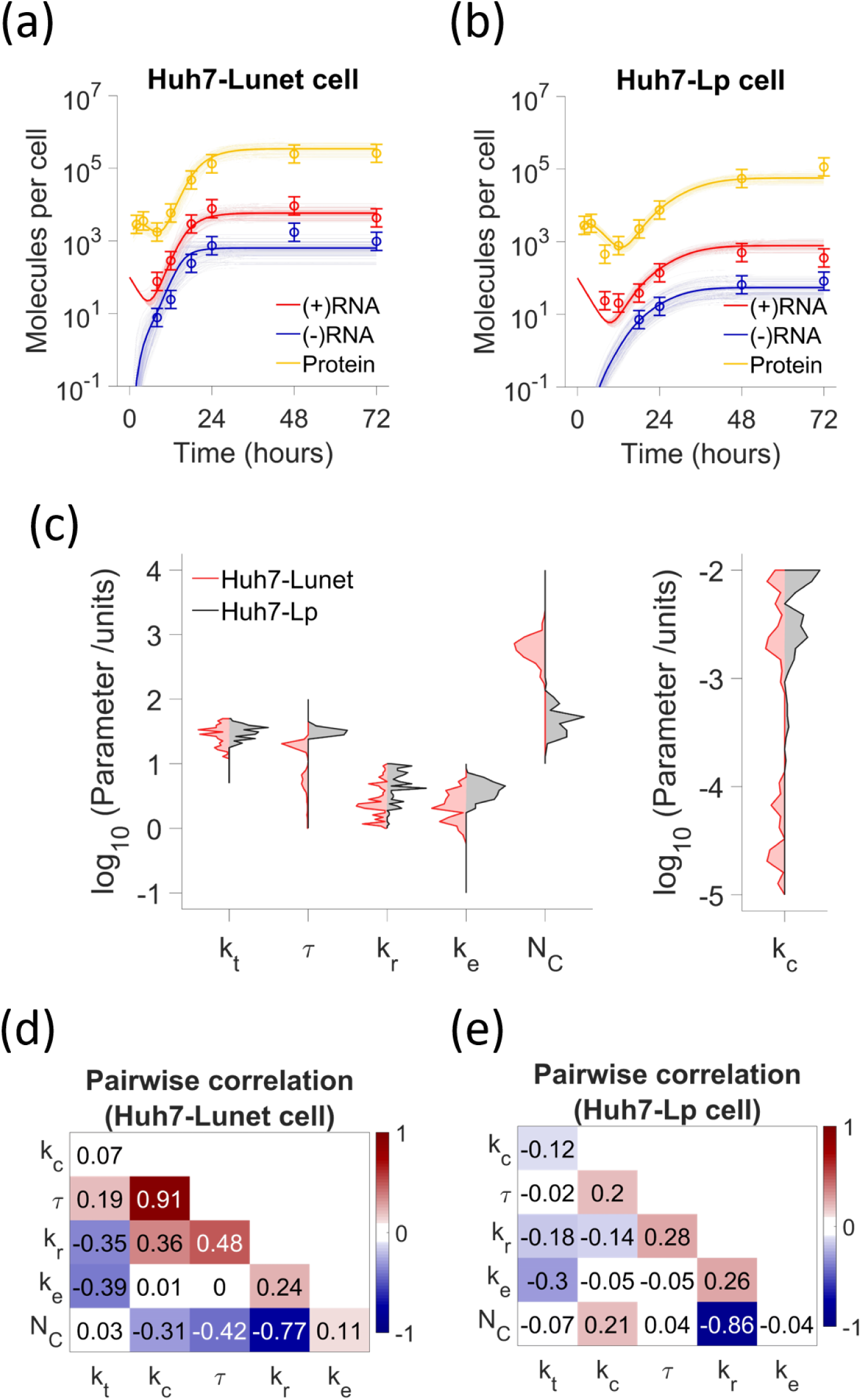
Parsing effects of host cell permissiveness. Model Fits (thin lines, best parameter fits; thick line, average of best fits) for the observed transfection dynamics of sgHCV (JFH1 strain) in (a) low permissive (Huh7-Lp) and (b) high permissive (Huh7-Lunet) cell lines. Initial condition used in both cases: *R*_*cyt*_ = 100. We consider constant relative error in observation. We ignore viral assembly (*k*_*a*_ = 0) for JFH1 strain. Data source: [4]. Hollow circles and error bars correspond to data used for fitting. (c) Comparison of parameter value distributions estimated for sgHCV transfection in Huh7-Lp (grey) and Huh7-Lunet (red). Pairwise correlation (see SI SM2) among the parameters estimated for sgHCV transfection in (d) Huh7-Lp and (e) Huh7-Lunet cell.

**Figure S3:**
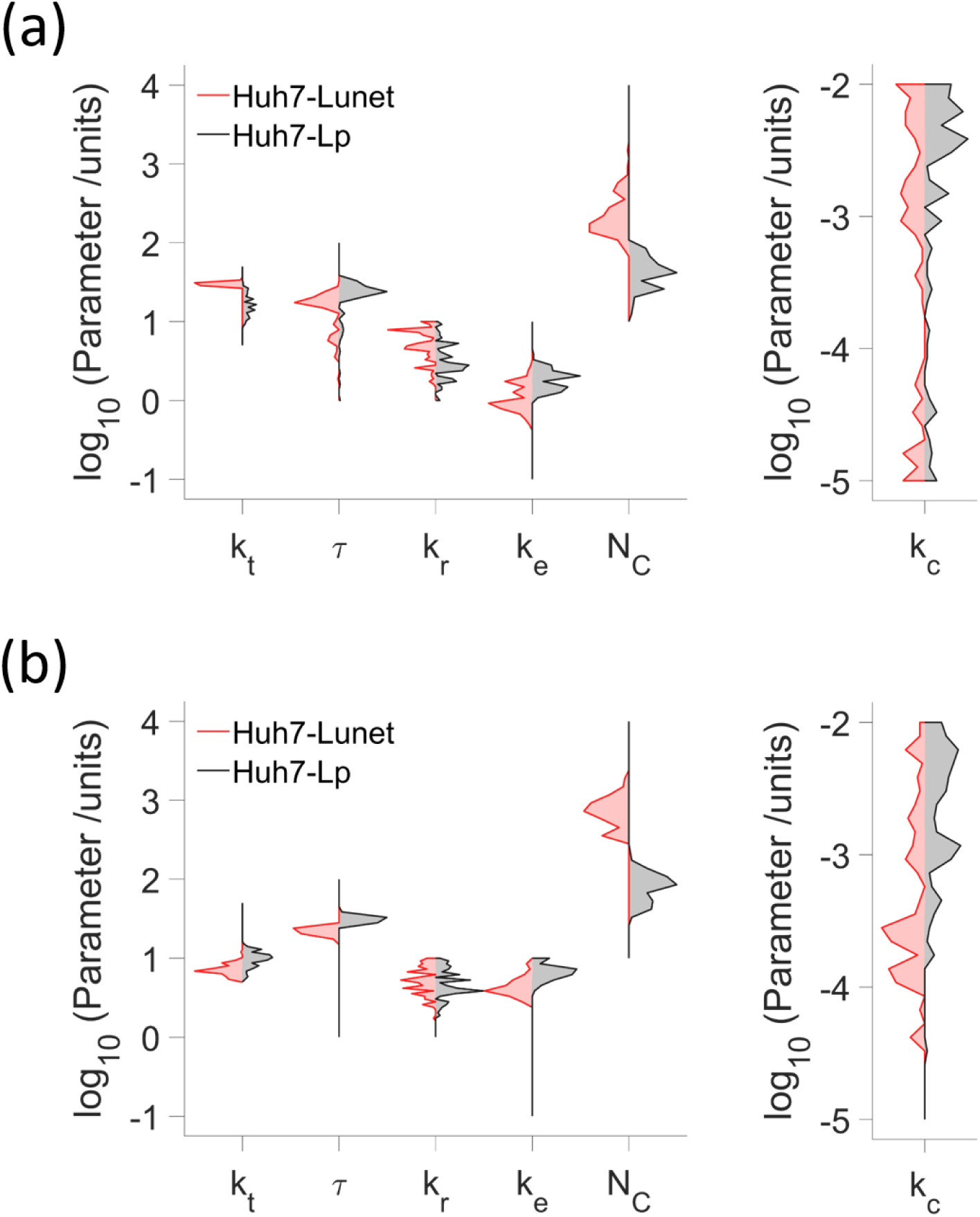
Fitting transfection dynamics using different initial conditions. Comparison of parameter value distributions estimated for sgHCV transfection in Huh7-Lp (grey) and Huh7-Lunet (red) using initial condition (a) *R*_*cyt*_ = 100 [using reported experimental error [4]] and (b) *R*_*cyt*_ = 1000 [using constant 25% relative error]. Data used for fitting and implementation details are identical to that corresponding to results shown in Figure S2.

**Figure S4:**
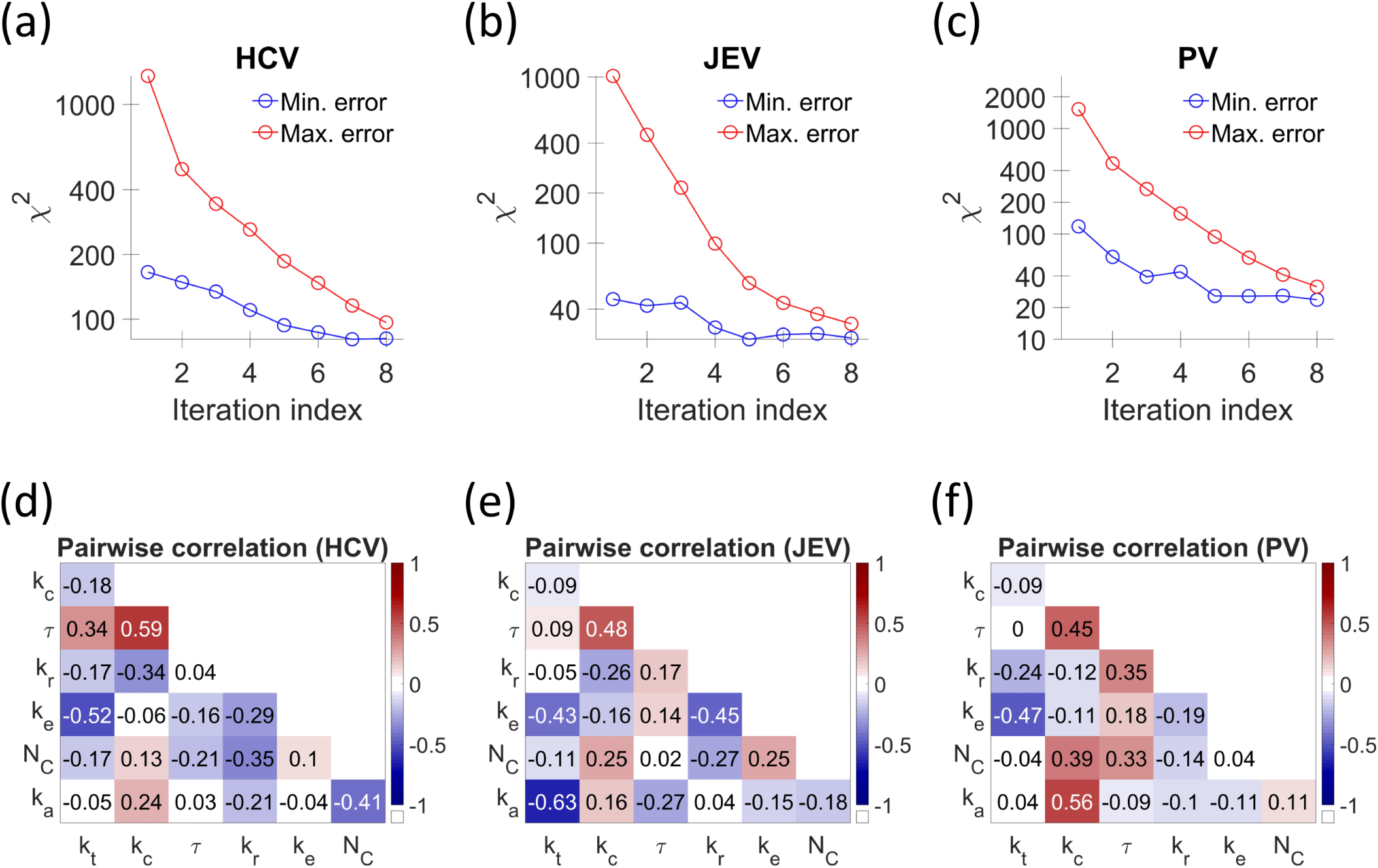
Algorithm convergence and practical identifiability analysis for fitting cellular dynamics of HCV, JEV and PV using 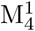. Variation of minimum (red) and maximum (blue) *χ*^2^, corresponding to parameter combinations selected, with each progressive iteration of iABC while fitting life cycle dynamics of (a) HCV, (b) JEV and (c) PV. Pairwise correlation (see SI SM2) in the values of parameters estimated for (d) HCV, (e) JEV and (f) PV.

**Figure S5:**
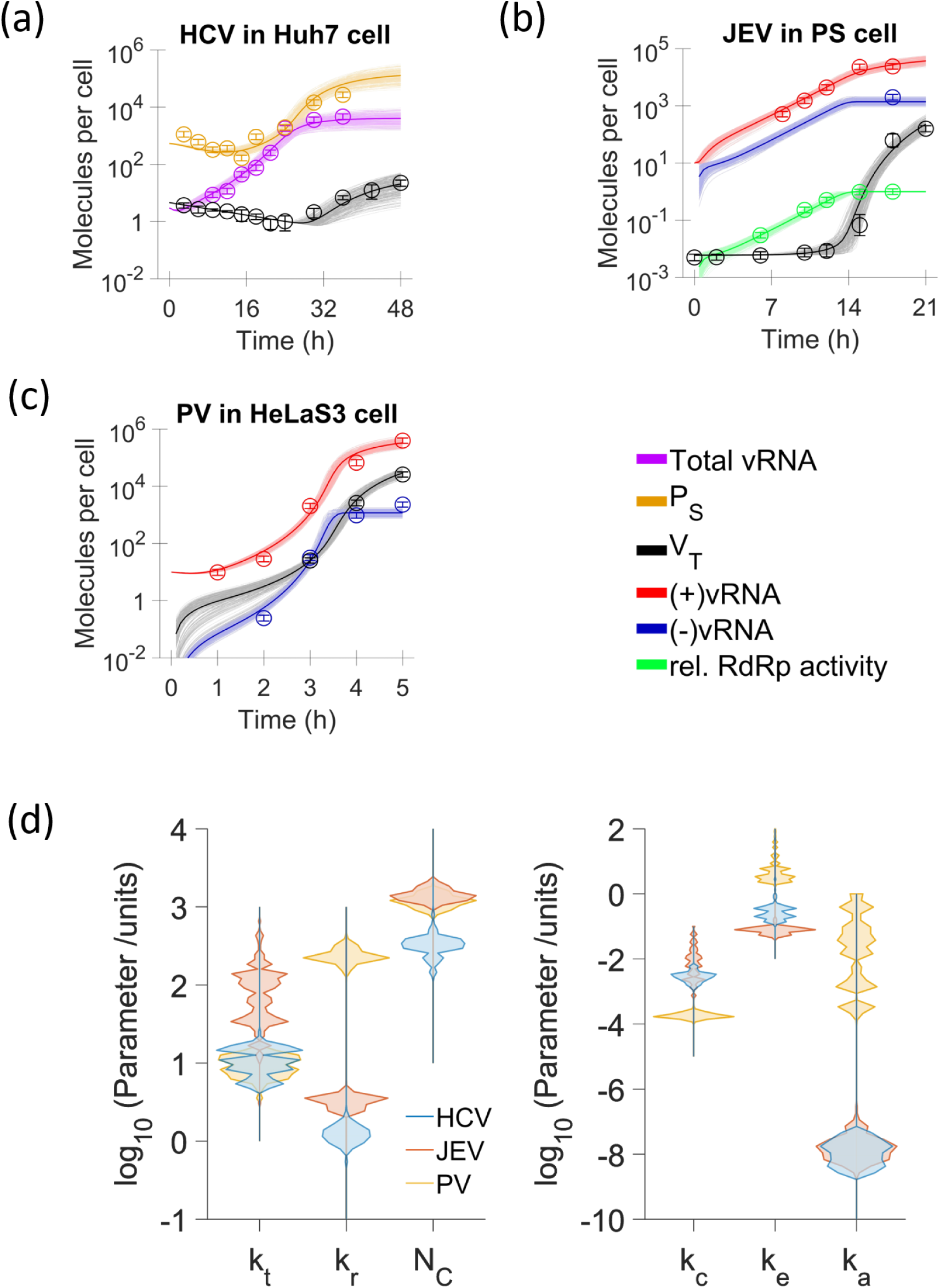
Analysing alternate model, M′: *f*_*CM*_ formulation variant where *f*_*CM*_ = 1. Model Fits (thin lines, best parameter fits; thick line, average of best fits) for the observed cellular life cycle dynamics of (a) HCV infection in Huh7 cells [1], (b) JEV infection in PS cells [32], and (c) PV infection in HeLaS3 cells [28]. (d) Comparison of parameter value distributions estimated for HCV, JEV and PV life cycle dynamics from iABC based model fitting (implementation details in SI SM3).

**Figure S6:**
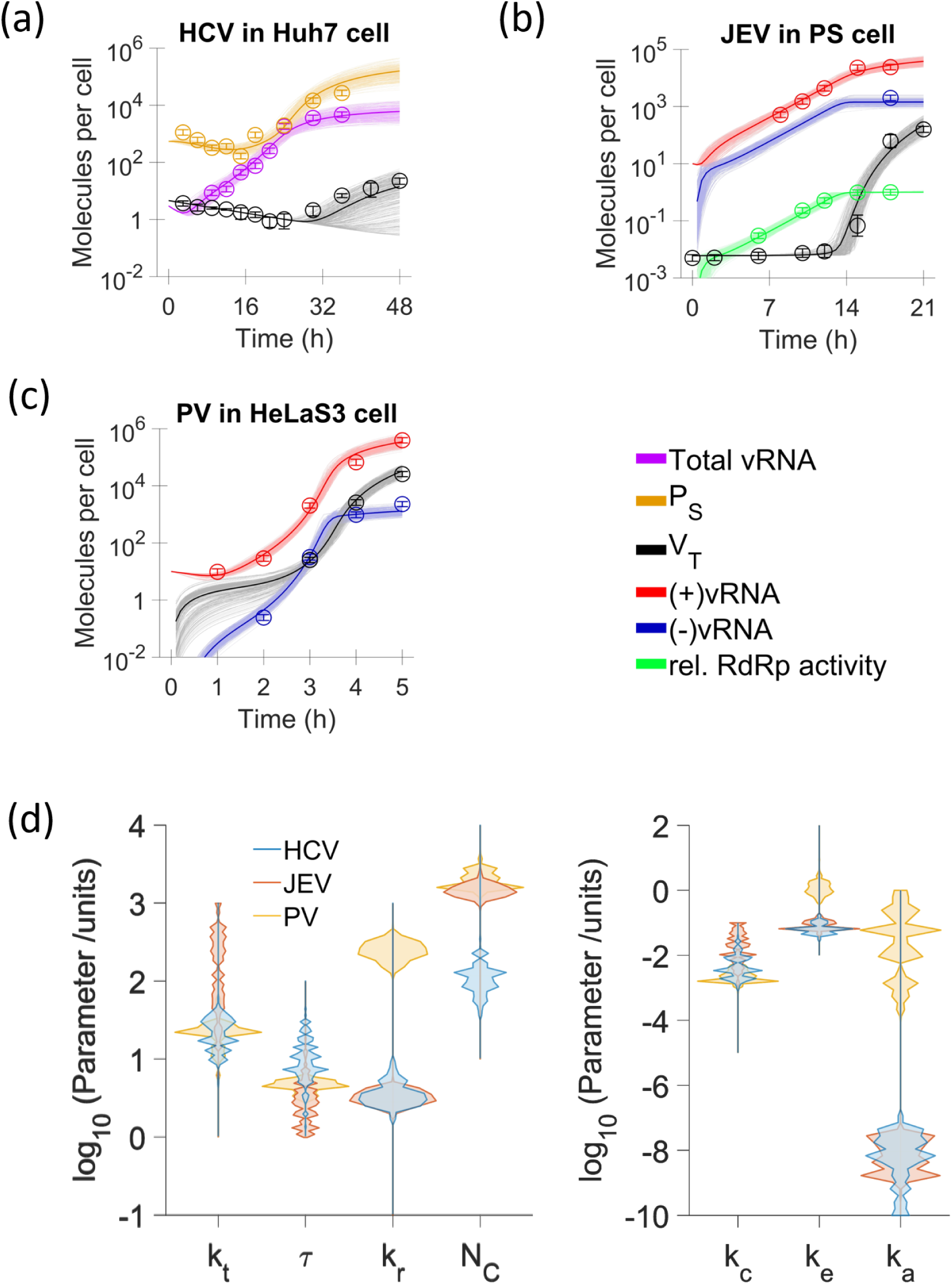
Analysing alternate model, 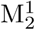: *f*_*CM*_ formulation variant where 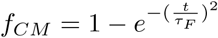. Model Fits (thin lines, best parameter fits; thick line, average of best fits) for the observed cellular life cycle dynamics of (a) HCV infection in Huh7 cells [1], (b) JEV infection in PS cells [32], and (c) PV infection in HeLaS3 cells [28]. (d) Comparison of parameter value distributions estimated for HCV, JEV and PV life cycle dynamics from iABC based model fitting (implementation details in SI SM3).

**Figure S7:**
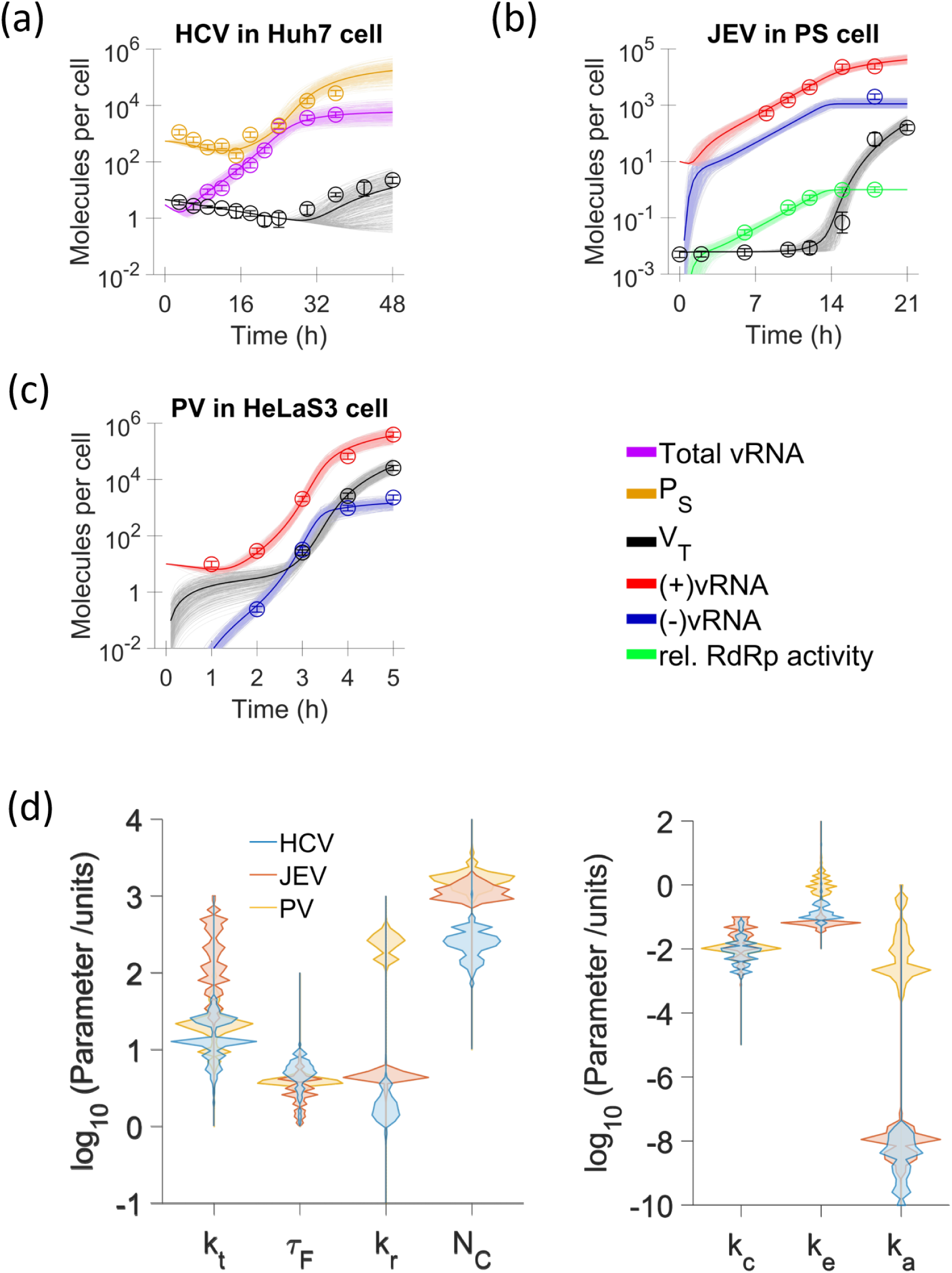
Analysing alternate model, 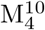: formulation assumes that 10 *P*_*NS*_ molecules are required to form one *RC*_*CM*_. Model Fits (thin lines, best parameter fits; thick line, average of best fits) for the observed cellular life cycle dynamics of (a) HCV infection in Huh7 cells [1], (b) JEV infection in PS cells [32], and (c) PV infection in HeLaS3 cells [28]. (d) Comparison of parameter value distributions estimated for HCV, JEV and PV life cycle dynamics from iABC based model fitting (implementation details in SI SM3).

**Figure S8:**
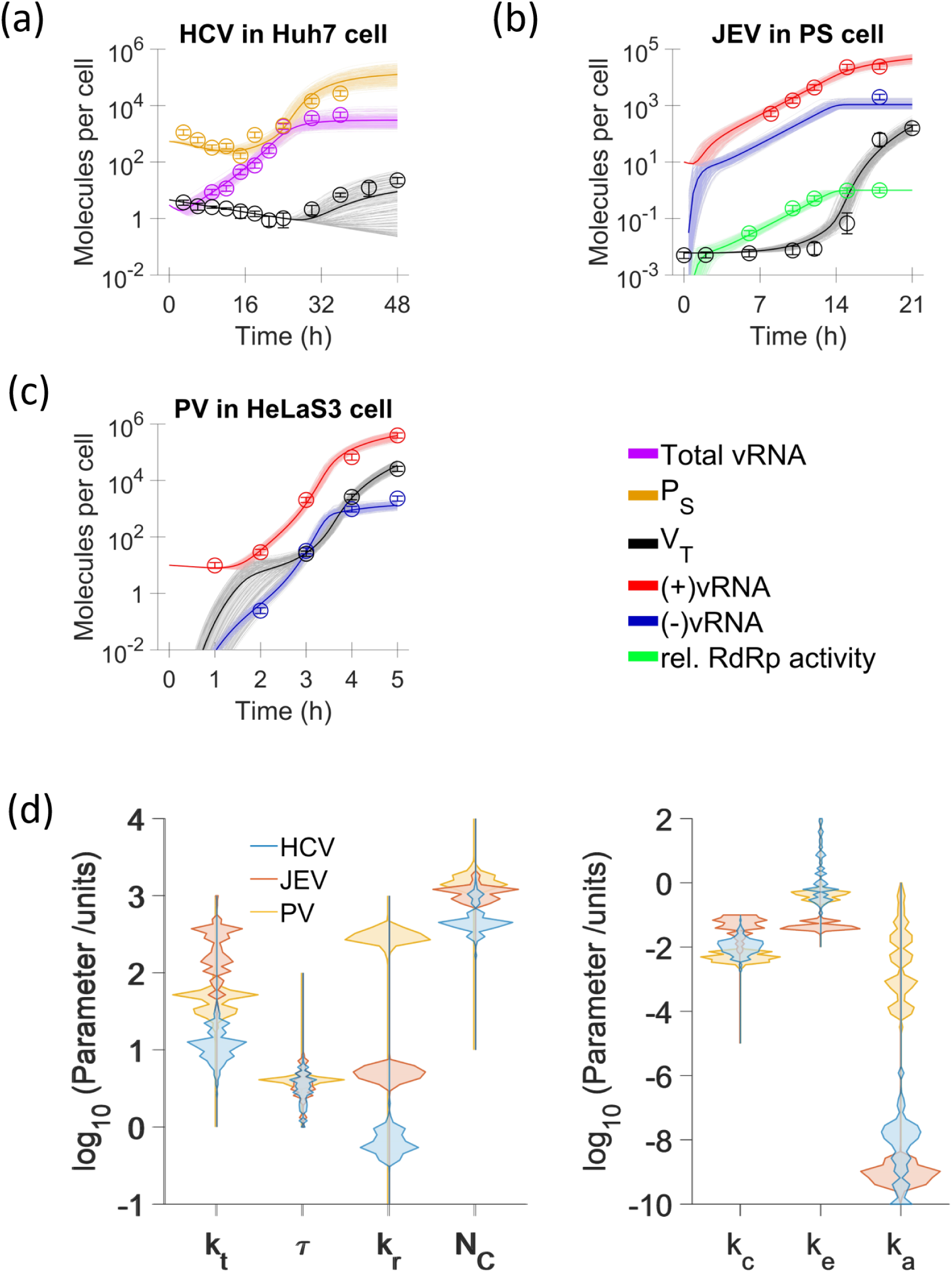
Analysing alternate model, W_4_: Packaging formulation variant where genomes from compartments associate with SP from cytoplasm to form new viral particles. Model Fits (thin lines, best parameter fits; thick line, average of best fits) for the observed cellular life cycle dynamics of (a) HCV infection in Huh7 cells [1], (b) JEV infection in PS cells [32], and (c) PV infection in HeLaS3 cells [28]. (d) Comparison of parameter value distributions estimated for HCV, JEV and PV life cycle dynamics from iABC based model fitting (implementation details in SI SM3).

**Figure S9:**
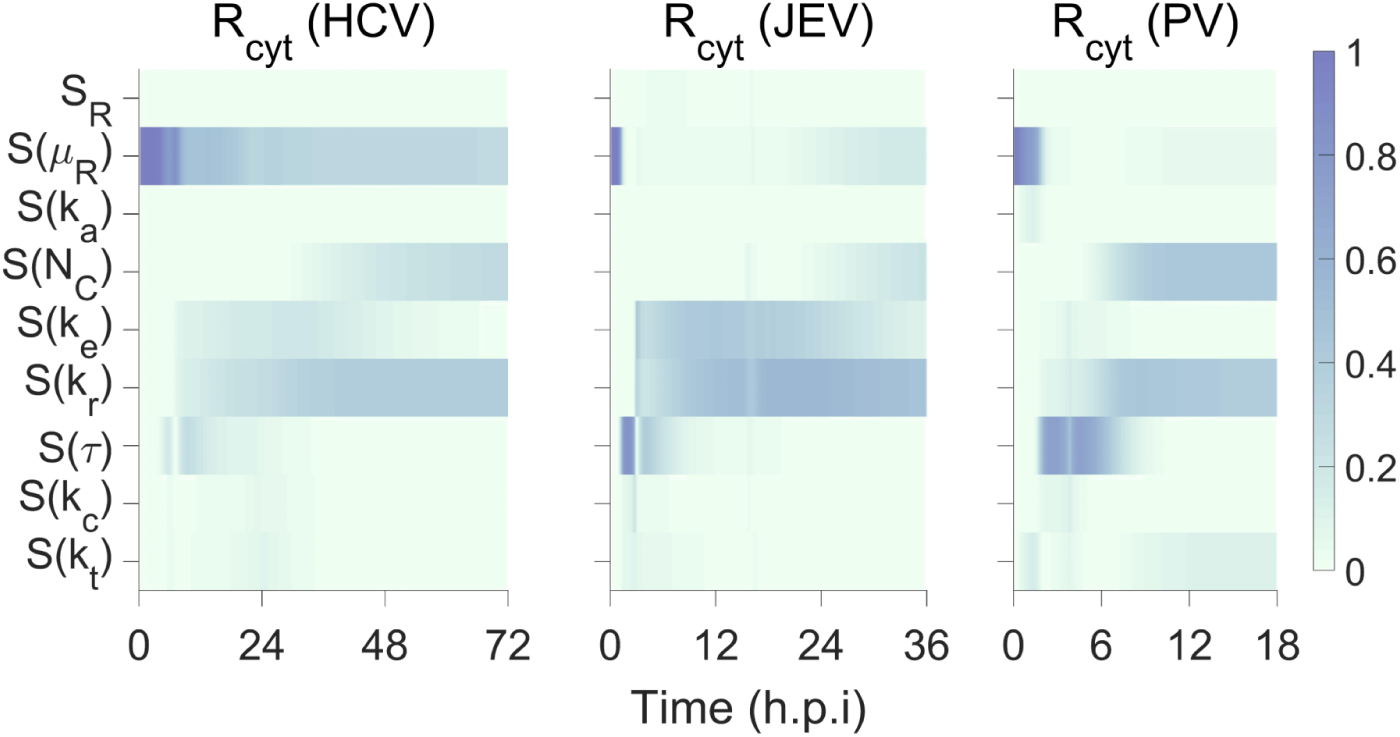
Parameter-temporal sensitivity profiles for dynamics of (+)vRNA in cytoplasm (*R*_*cyt*_) for HCV, JEV and PV (at seeding *R*_*cyt*_(*t* = 0)=3) are shown. Here *S*(*X*) denote the profile associated with parameter, X, and *S*_*R*_ = *S*(*µ*_*P*_) + *S*(*µ*_*V*_) + *S*(*dummy*). Time axis is not to scale across profiles for different viruses.

**Figure S10:**
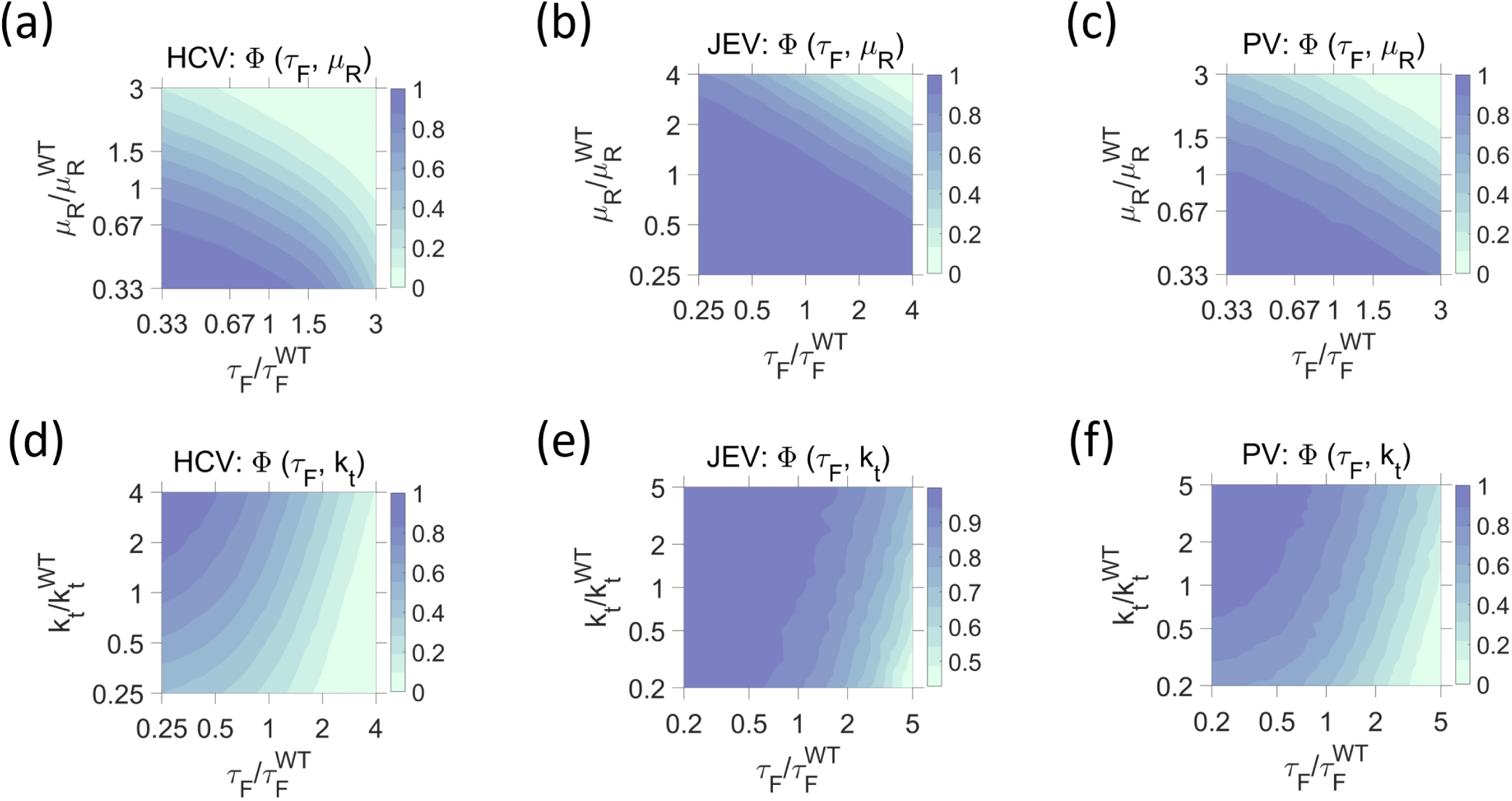
Cellular infectivity profiles. Φ(*τ*_*F*_, *µ*_*R*_) evaluates the cellular infectivity at different values of compartment formation delay (*τ*_*F*_) and degradation rate of viral +RNA in cytoplasm (*µ*_*R*_) for (a) JEV and (b) PV at seeding, N = 3. Φ(*τ*_*F*_, *k*_*t*_) evaluates the cellular infectivity at different values of *τ*_*F*_ and viral translation rate (*k*_*t*_) for (d) HCV, (e) JEV and (f) PV at seeding, N = 3.

**Figure S11:**
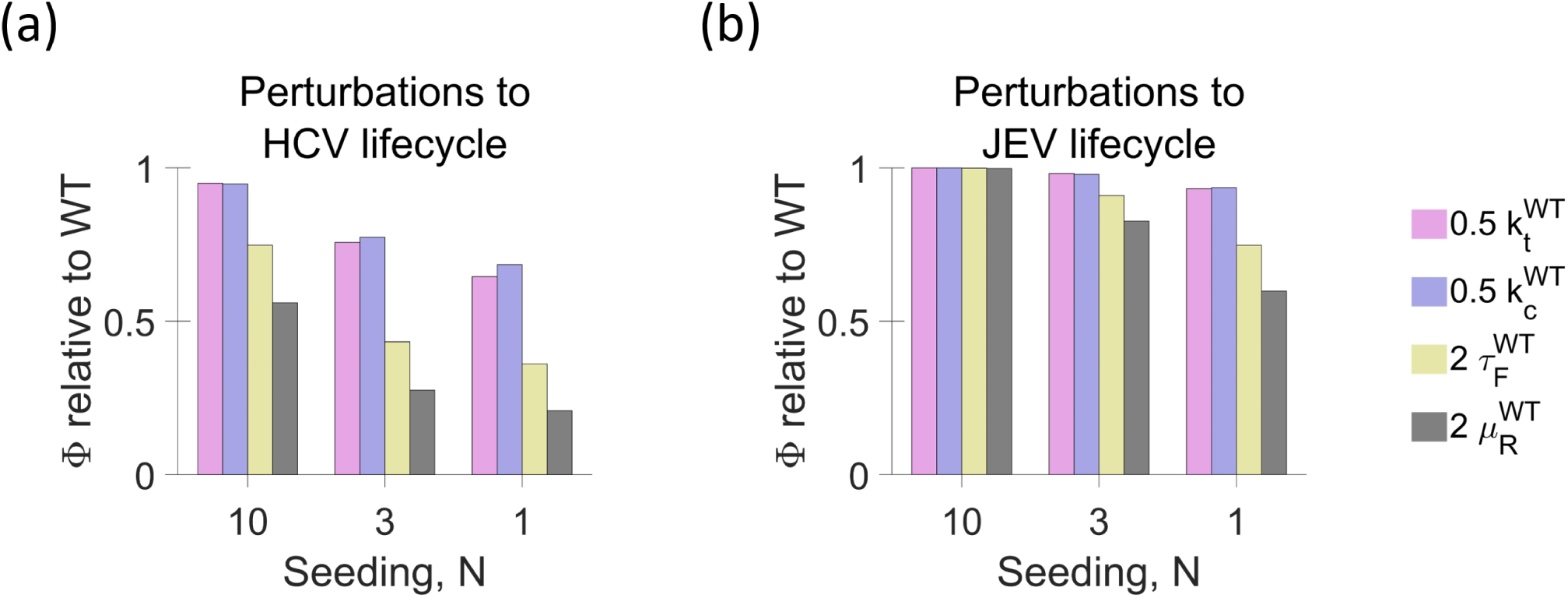
Seeding dependent fold change in Φ due to change in life cycle parameter value for (a) HCV and (b) JEV. 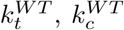 and 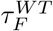 correspond to estimates for correspond to the estimates for the respective virus (Table 1, Main text), and 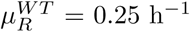.

**Figure S12:**
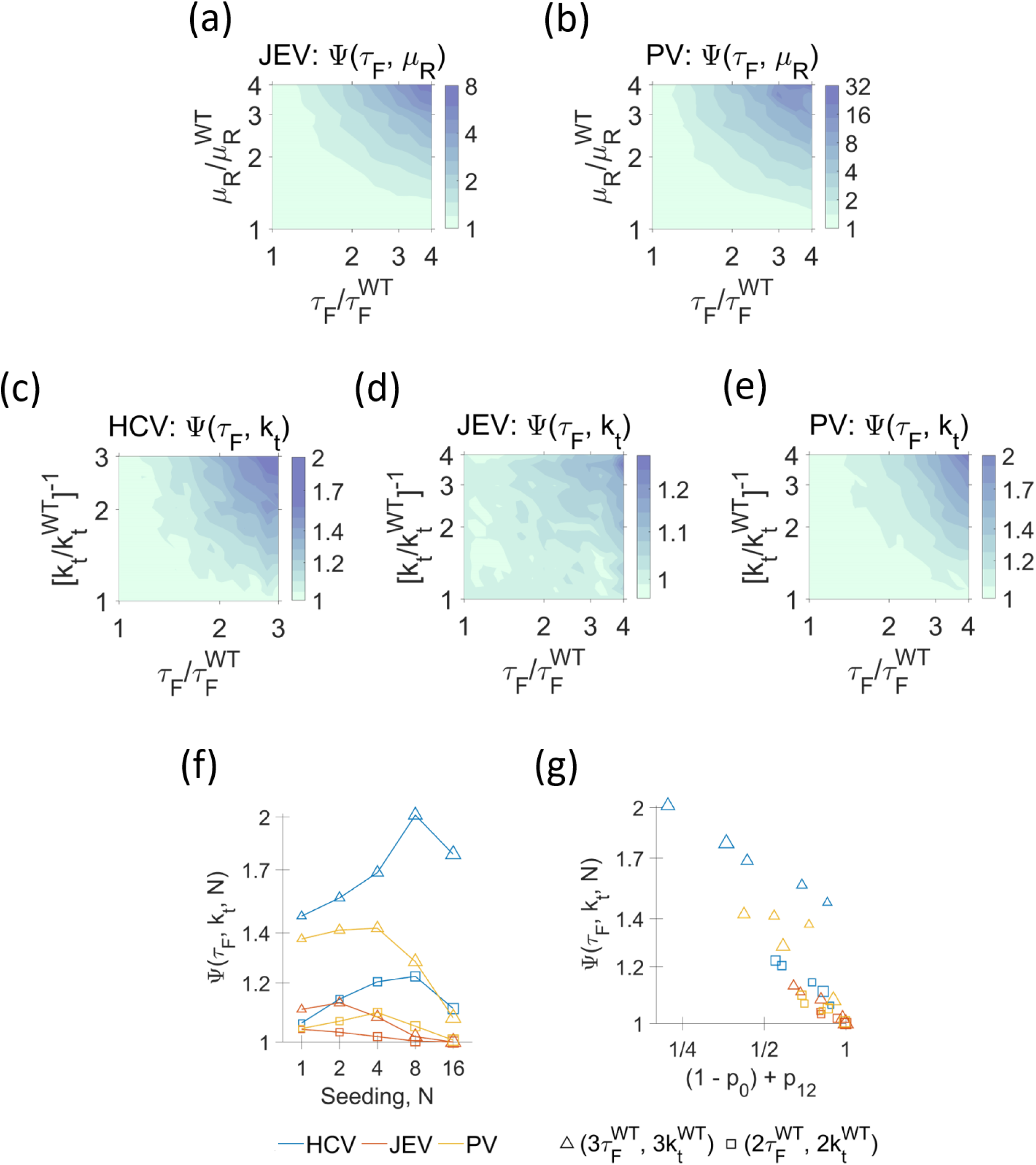
Synergestic strategies to reduce cellular infectivity. Ψ(*τ*_*F*_, *µ*_*R*_) evaluates the Bliss synergy between compartment formation delay (*τ*_*F*_) and degradation rate of viral +RNA in cytoplasm (*µ*_*R*_) for (a) JEV and (b) PV at seeding, N = 3. Ψ(*τ*_*F*_, *k*_*t*_) evaluates the Bliss synergy between *τ*_*F*_ and viral translation rate (*k*_*t*_) for (c) HCV, (d) JEV and (e) PV at seeding, N = 3.Color scale may vary across the sub-figures (a-e). (f) Variation of Ψ(*τ*_*F*_, *µ*_*R*_) for various levels of change in parameter values (denoted by different markers) with viral seeding (N) for HCV (blue), JEV (red) and PV (yellow). (g) Ψ(*τ*_*F*_, *k*_*t*_) shows a negative correlation with {(1 − *p*_0_) + *p*_12_}. 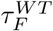 and 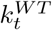 correspond to the estimates for the respective virus (Table 1, Main text), and 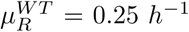. Marker properties are same in (f) and (g) and larger markers correspond to higher N.

**Table S1:** Compartment formation dynamics: Parameter estimation and goodness of fit*

| Infection system | Estimated $\tau_S$ (in h) | | | | Estimated $N_{CM}$ | | | | $\chi^2$ | | | | Data source |
| --- | --- | --- | --- | --- | --- | --- | --- | --- | --- | --- | --- | --- | --- |
|  | n=2 | n=3 | n=4 | n=5 | n=2 | n=3 | n=4 | n=5 | n=2 | n=3 | n=4 | n=5 |  |
| 1. MHV | 5.9 | 5.3 | 4.8 | 4.7 | 17.8 | 17.8 | 17.8 | 17.8 | 13.6 | 7.3 | 5.7 | 5.4 | [33] |
| 2. EAV | 6.7 | 6 | 5.7 | 5.3 | 223.9 | 223.9 | 219.3 | 177.8 | 29.2 | 11.8 | 2.8 | 0.2 | [16] |
| 3. ZIKV<sup>M</sup> | 20.1 | 15 | 13.3 | 13.3 | 70.8 | 57.1 | 56.2 | 56.2 | 0.5 | 2.6 | 6.1 | 9.3 | [8] |
| 4. ZIKV<sup>H</sup> | 37.6 | 26.6 | 23.7 | 21.1 | 71.1 | 70.8 | 70.8 | 70.8 | 12.2 | 2.1 | 1.3 | 1.5 | [8] |
| 5. HCV | 37.6 | 26.6 | 23.7 | 21.1 | 446.7 | 354.8 | 354.8 | 354.8 | 15.5 | 8.3 | 3.6 | 1 | [27] |

MHV: Mouse Hepatitis virus infection in HeLaS3 cells; EAV: Equine Arteritis virus in Vero E6 cells; ZIKV<sup>M</sup>: Zika viral strain MR766 infection in Huh cells; ZIKV<sup>H</sup>: Zika viral strain H/PF/2013 infection in Huh cells; HCV: Hepatitis C virus infection in Huh cells.
The estimates correspond to median of the estimated distribution obtained in the fifth iteration of iABC.
We fit dynamics of compartments formed, $X$ , using eq. S1 for $n = 2, 3$ and $4$ .
where $N_{CM}$ is the carrying capacity with respect to replication compartments in the infected host cell using $f_{CM}$ given by eq. 1 (main text). We fit $\log_{10}(\tau_S$ [in h]) and $\log_{10}(N_{CM})$ for each virus using initial guess for parameter value distribution as $u(0, 2)$ and $u(0, 3)$ respectively. Here $u(a, b)$ represent the uniform distribution with $a$ and $b$ as the lower and upper bounds of the distribution respectively.

**Table S2:** Cellular life cycle model*: Parameter estimation summary

| Parameter<br>(Unit) | Estimated value: $Q_2, (Q_1, Q_3)$ # | | |
| --- | --- | --- | --- |
|  | HCV in Huh7 cell | JEV in PS cells | PV in HeLaS3 cells |
| $k_t$<br>($h^{-1}$) | 23.7<br>(21.3, 26.1) | $1.6 \times 10^2$<br>(1.2, 1.9) $\times 10^2$ | 18.9<br>(17.4, 20.6) |
| $k_c$<br>(molecules$^{-1}$.h$^{-1}$) | $2.6 \times 10^{-3}$<br>(2, 3.4) $\times 10^{-3}$ | $1.6 \times 10^{-2}$<br>(0.8, 3.1) $\times 10^{-2}$ | $1.2 \times 10^{-2}$<br>(1, 1.4) $\times 10^{-2}$ |
| $\tau_F$<br>(h) | 5.8<br>(5.2, 6.3) | 2.6<br>(1.8, 2.9) | 4.2<br>(4.1, 4.3) |
| $k_r$<br>($h^{-1}$) | 3.6<br>(3.4, 3.9) | 3.7<br>(3.5, 3.9) | $2.2 \times 10^2$<br>(2, 2.4) $\times 10^2$ |
| $k_e$<br>($h^{-1}$) | $6.6 \times 10^{-2}$<br>(5.9, 7.2) $\times 10^{-2}$ | $7.2 \times 10^{-2}$<br>(6.7, 8) $\times 10^{-2}$ | 1.1<br>(1, 1.2) |
| $N_C$<br>(molecules) | $8.8 \times 10^1$<br>(8.1, 9.8) $\times 10^1$ | $1.21 \times 10^3$<br>(1.15, 1.31) $\times 10^3$ | $1.62 \times 10^3$<br>(1.53, 1.78) $\times 10^3$ |
| $k_a$<br>(molecules$^{-1}$.h$^{-1}$) | $3.6 \times 10^{-8}$<br>(2.9, 4.7) $\times 10^{-8}$ | $8 \times 10^{-9}$<br>(6.4, 10.2) $\times 10^{-9}$ | $5.6 \times 10^{-3}$<br>(3.2, 20.3) $\times 10^{-3}$ |

\* Using life cycle model given in main text ( $M_4^1$ ).

**Table S3:** Comparing parameter estimation for sgHCV (JFH1 strain) life cycle in host cells of different permissivity

| Parameter<br>(Unit) | Estimated value: $Q_2, (Q_1, Q_3)$ <sup>#</sup> | |
| --- | --- | --- |
|  | JFH1 in Huh7-Lp cell | JFH1 in Huh7-Lunet cell |
| $k_t$<br>(h<sup>-1</sup>) | 30.7<br>(25.2, 35.8) | 28.3<br>(21, 34.4) |
| $k_c$<br>(molecules<sup>-1</sup>.h<sup>-1</sup>) | $3.5 \times 10^{-3}$<br>(2.1, 7.5) $\times 10^{-3}$ | $1.6 \times 10^{-3}$<br>(3.4) $\times 10^{-3}$ |
| $\tau_F$<br>(h) | 29.9<br>(27.5, 32.5) | 16.4<br>(5.4, 19.6) |
| $k_r$<br>(h<sup>-1</sup>) | 4.8<br>(3.3, 7) | 2.4<br>(1.7, 3.7) |
| $k_e$<br>(h<sup>-1</sup>) | 3.7<br>(3, 4.6) | 2.1<br>(1.4, 3) |
| $N_C$<br>(molecules) | 51.2<br>(34.2, 61.9) | 603.1<br>(455.3, 768.8) |

<sup>#</sup> $Q_1, Q_2$ and $Q_3$ represent the first, second (median) and third quartiles respectively of the estimated distribution (after final iteration of iABC). For the analysis we assume that 100 viral genomes enter each cell. See section SI SM3 for further implementation details. Data used for fitting was taken from [4].

**Table S4:**
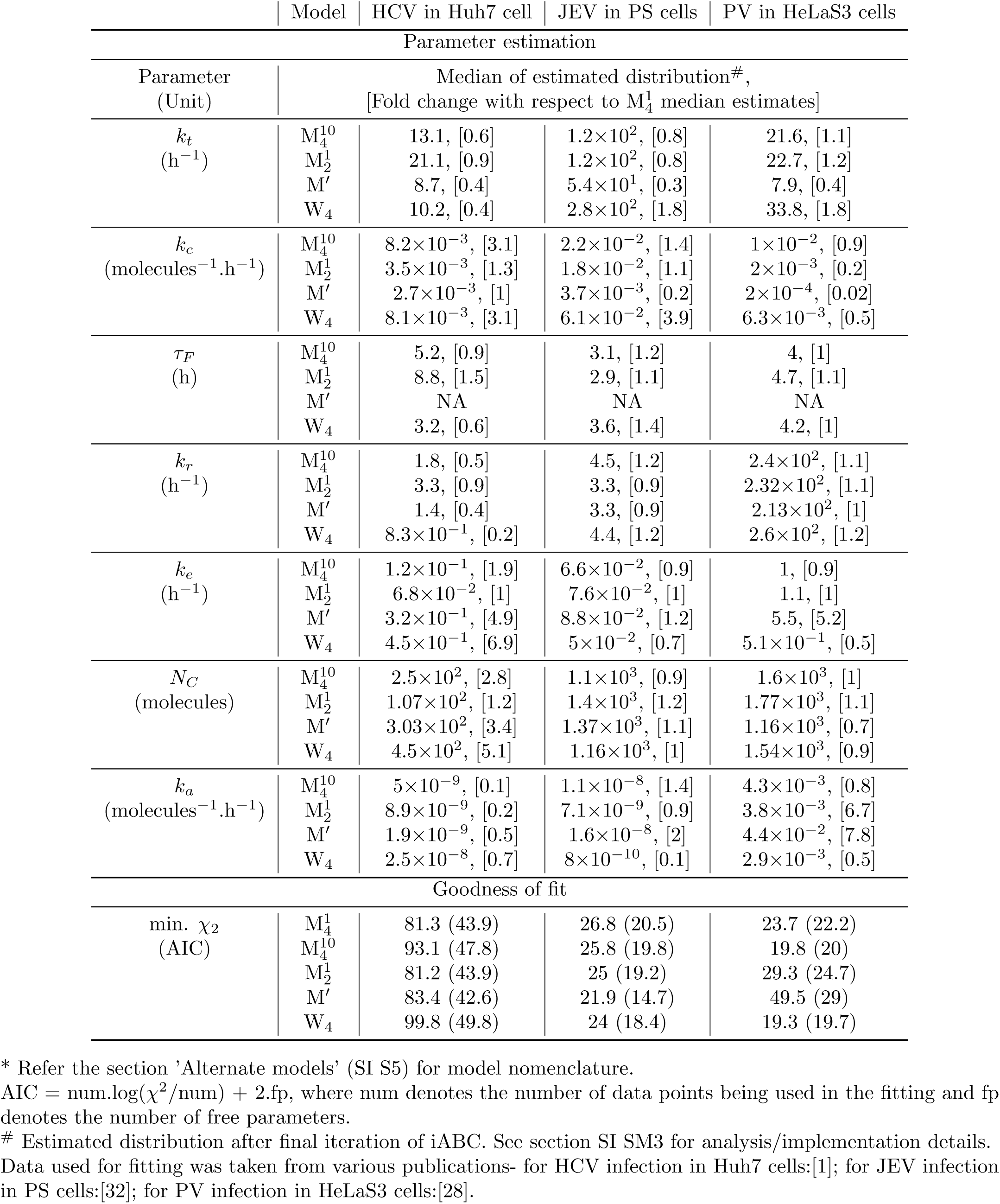
Comparing parameter estimation and goodness of fit for model variants

## References

[1] S. Miller and J. Krijnse-Locker. Modification of intracellular membrane structures for virus repli-cation. Nat. Rev. Microbiol., 6(5):363–374, May 2008.

[2] J. A. den Boon and P. Ahlquist. Organelle-like membrane compartmentalization of positive-strand RNA virus replication factories. Annu. Rev. Microbiol., 64:241–256, 2010.

[3] D. Paul and R. Bartenschlager. Architecture and biogenesis of plus-strand RNA virus replication factories. World J Virol, 2(2):32–48, May 2013.

[4] I. Romero-Brey and R. Bartenschlager. Membranous replication factories induced by plus-strand RNA viruses. Viruses, 6(7):2826–2857, Jul 2014.

[5] M. J. Wu, P. Y. Ke, J. T. Hsu, C. T. Yeh, and J. T. Horng. Reticulon 3 interacts with NS4B of the hepatitis C virus and negatively regulates viral replication by disrupting NS4B self-interaction. Cell. Microbiol., 16(11):1603–1618, Nov 2014.

[6] W. F. Tang, S. Y. Yang, B. W. Wu, J. R. Jheng, Y. L. Chen, C. H. Shih, K. H. Lin, H. C. Lai, P. Tang, and J. T. Horng. Reticulon 3 Binds the 2C Protein of Enterovirus 71 and Is Required for Viral Replication. J. Biol. Chem., 282(8):5888–5898, Feb 2007.

[7] T. E. Aktepe, S. Liebscher, J. E. Prier, C. P. Simmons, and J. M. Mackenzie. The Host Protein Reticulon 3.1A Is Utilized by Flaviviruses to Facilitate Membrane Remodelling. Cell Rep, 21(6):1639–1654, Nov 2017.

[8] Anna Lundin, Ronald Dijkman, Tomas Bergström, Nina Kann, Beata Adamiak, and Hannoun. Targeting membrane-bound viral RNA synthesis reveals potent inhibition of diverse coronaviruses including the middle East respiratory syndrome virus. PLoS Pathog., 10(5):e1004166, May 2014.

[9] Obdulio García-Nicolás, Philip V’kovski, Nathalie J Vielle, Nadine Ebert, Roland Züst, Jasmine Portmann, Hanspeter Stalder, Véronique Gaschen, Gabrielle Vieyres, Michael Stoffel, et al. The Small-Compound Inhibitor K22 Displays Broad Antiviral Activity against Different Members of the Family Flaviviridae and Offers Potential as a Panviral Inhibitor. Antimicrob. Agents Chemother., 62(11), 11 2018.

[10] C. Berger, I. Romero-Brey, D. Radujkovic, R. Terreux, M. Zayas, D. Paul, C. Harak, S. Hoppe, M. Gao, F. Penin, V. Lohmann, and R. Bartenschlager. Daclatasvir-like inhibitors of NS5A block early biogenesis of hepatitis C virus-induced membranous replication factories, independent of RNA replication. Gastroenterology, 147(5):1094–1105, Nov 2014.

[11] K. L. Berger, S. M. Kelly, T. X. Jordan, M. A. Tartell, and G. Randall. Hepatitis C Virus Stimulates the Phosphatidylinositol 4-Kinase III Alpha-Dependent Phosphatidylinositol 4-Phosphate Production That Is Essential for Its Replication. J. Virol., 85(17):8870–8883, Sep 2011.

[12] M. B. Schulte and R. Andino. Single-cell analysis uncovers extensive biological noise in poliovirus replication. J. Virol., 88(11):6205–6212, Jun 2014.

[13] F. S. Heldt, S. Y. Kupke, S. Dorl, U. Reichl, and T. Frensing. Single-cell analysis and stochastic modelling unveil large cell-to-cell variability in influenza A virus infection. Nat Commun, 6:8938, Nov 2015.

[14] R. Srivastava, L. You, J. Summers, and J. Yin. Stochastic vs. deterministic modeling of intracellular viral kinetics. J. Theor. Biol., 218(3):309–321, Oct 2002.

[15] B. Snijder, R. Sacher, P. Rämö, E. M. Damm, P. Liberali, and L. Pelkmans. Population context determines cell-to-cell variability in endocytosis and virus infection. Nature, 461(7263):520–523, Sep 2009.

[16] H. Dahari, R. M. Ribeiro, C. M. Rice, and A. S. Perelson. Mathematical Modeling of Subgenomic Hepatitis C Virus Replication in Huh-7 Cells. J. Virol., 81(2):750–760, Jan 2007.

[17] Marco Binder, Nurgazy Sulaimanov, Diana Clausznitzer, Manuel Schulze, Christian M Hüber, Simon M Lenz, Johannes P Schlöder, Martin Trippler, Ralf Bartenschlager, Volker Lohmann, et al. Replication Vesicles are Load-and Choke-Points in the Hepatitis C Virus Lifecycle. PLoS Pathog., 9(8), 2013.

[18] R. R. Regoes, S. Crotty, R. Antia, and M. M. Tanaka. Optimal replication of poliovirus within cells. Am. Nat., 165(3):364–373, Mar 2005.

[19] M. B. Schulte, J. A. Draghi, J. B. Plotkin, and R. Andino. Experimentally guided models reveal replication principles that shape the mutation distribution of RNA viruses. Elife, 4, Jan 2015.

[20] A. K. McLean, F. Luciani, and M. M. Tanaka. Trade-offs in resource allocation in the intracellular life-cycle of hepatitis C virus. J. Theor. Biol., 267(4):565–572, Dec 2010.

[21] D. Clausznitzer, J. Harnisch, and L. Kaderali. Multi-scale model for hepatitis C viral load kinetics under treatment with direct acting antivirals. Virus Res., 218:96–101, 06 2016.

[22] T. R. Aunins, K. A. Marsh, G. Subramanya, S. L. Uprichard, A. S. Perelson, and A. Chatterjee. Intracellular Hepatitis C Virus Modeling Predicts Infection Dynamics and Viral Protein Mechanisms. J. Virol., 92(11), 06 2018.

[23] C. Zitzmann, B. Schmid, A. Ruggieri, A. S. Perelson, M. Binder, R. Bartenschlager, and L. Kaderali. A Coupled Mathematical Model of the Intracellular Replication of Dengue Virus and the Host Cell Immune Response to Infection. Front Microbiol, 11:725, 2020.

[24] Ashley I Teufel, Wu Liu, Jeremy A Draghi, Craig E Cameron, and Claus O Wilke. Uncovering modeling features of viral replication dynamics from high-throughput single-cell virology experiments. bioRxiv, 2020.

[25] J. Yin and J. Redovich. Kinetic Modeling of Virus Growth in Cells. Microbiol. Mol. Biol. Rev., 82(2), 06 2018.

[26] S. Miyashita, K. Ishibashi, H. Kishino, and M. Ishikawa. Viruses roll the dice: the stochastic behavior of viral genome molecules accelerates viral adaptation at the cell and tissue levels. PLoS Biol., 13(3):e1002094, Mar 2015.

[27] Carolin Zitzmann and Lars Kaderali. Mathematical analysis of viral replication dynamics and antiviral treatment strategies: from basic models to age-based multi-scale modeling. Frontiers in microbiology, 9:1546, 2018.

[28] A. S. Perelson, E. Herrmann, F. Micol, and S. Zeuzem. New kinetic models for the hepatitis C virus. Hepatology, 42(4):749–754, Oct 2005.

[29] T. Nguyen and J. Guedj. HCV Kinetic Models and Their Implications in Drug Development. CPT Pharmacometrics Syst Pharmacol, 4(4):231–242, Apr 2015.

[30] A. Zimmer, I. Katzir, E. Dekel, A. E. Mayo, and U. Alon. Prediction of multidimensional drug dose responses based on measurements of drug pairs. Proc. Natl. Acad. Sci. U.S.A., 113(37):10442–10447, 09 2016.

[31] P. Padmanabhan and N. M. Dixit. Modeling Suggests a Mechanism of Synergy Between Hepatitis C Virus Entry Inhibitors and Drugs of Other Classes. CPT Pharmacometrics Syst Pharmacol, 4(8):445–453, Aug 2015.

[32] P. D. Uchil and V. Satchidanandam. Characterization of RNA synthesis, replication mechanism, and in vitro RNA-dependent RNA polymerase activity of Japanese encephalitis virus. Virology, 307(2):358–371, Mar 2003.

[33] Ronald WAL Limpens, Hilde M van der Schaar, Darshan Kumar, Abraham J Koster, Eric J Snijder, Frank JM van Kuppeveld, and Montserrat Bárcena. The Transformation of Enterovirus Replication Structures: a Three-Dimensional Study of Single- and Double-Membrane Compartments. mBio, 2(5), 2011.

[34] Barbara van der Hoeven, Diede Oudshoorn, Abraham J Koster, Eric J Snijder, Marjolein Kikkert, and Montserrat Bárcena. Biogenesis and architecture of arterivirus replication organelles. Virus Res., 220:70–90, 07 2016.

[35] David Paul, Inés Romero-Brey, Jérôme Gouttenoire, Savina Stoitsova, Jacomine Krijnse-Locker, Darius Moradpour, and Ralf Bartenschlager. NS4B self-interaction through conserved C-terminal elements is required for the establishment of functional hepatitis C virus replication complexes. “J. Virol”., 85(14):6963–6976, 2011.

[36] Terence L Wagner, Hsin-I Wu, Peter JH Sharpe, and RN Coulson. Modeling distributions of insect development time: a literature review and application of the Weibull function. Annals of the Entomological Society of America, 77(5):475–483, 1984.

[37] Kévin Knoops, Montserrat Bárcena, Ronald WAL Limpens, Abraham J Koster, A Mieke Mommaas, and Eric J Snijder. Ultrastructural Characterization of Arterivirus Replication Structures: Reshap-ing the Endoplasmic Reticulum To Accommodate Viral RNA Synthesis. J. Virol., 86(5):2474–2487, Mar 2012.

[38] M. Ulasli, M. H. Verheije, C. A. de Haan, and F. Reggiori. Qualitative and quantitative ultrastructural analysis of the membrane rearrangements induced by coronavirus. Cell. Microbiol., 12(6):844–861, Jun 2010.

[39] I. Romero-Brey, A. Merz, A. Chiramel, J. Y. Lee, P. Chlanda, U. Haselman, R. Santarella-Mellwig, A. Habermann, S. Hoppe, S. Kallis, P. Walther, C. Antony, J. Krijnse-Locker, and R. Bartenschlager. Three-Dimensional Architecture and Biogenesis of Membrane Structures Associated with Hepatitis C Virus Replication. PLoS Pathog., 8(12):e1003056, 2012.

[40] M. Cortese, S. Goellner, E. G. Acosta, C. J. Neufeldt, O. Oleksiuk, M. Lampe, U. Haselmann, C. Funaya, N. Schieber, P. Ronchi, M. Schorb, P. Pruunsild, Y. Schwab, L. Chatel-Chaix, A. Ruggieri, and R. Bartenschlager. Ultrastructural Characterization of Zika Virus Replication Factories. Cell Rep, 18(9):2113–2123, 02 2017.

[41] M. Münster, A. P laszczyca, M. Cortese, C. J. Neufeldt, S. Goellner, G. Long, and R. Bartenschlager. A Reverse Genetics System for Zika Virus Based on a Simple Molecular Cloning Strategy. Viruses, 10(7), 07 2018.

[42] B. Brandenburg, L. Y. Lee, M. Lakadamyali, M. J. Rust, X. Zhuang, and J. M. Hogle. Imaging poliovirus entry in live cells. PLoS Biol., 5(7):e183, Jul 2007.

[43] J. W. Oh, T. Ito, and M. M. Lai. A recombinant hepatitis C virus RNA-dependent RNA polymerase capable of copying the full-length viral RNA. J. Virol., 73(9):7694–7702, Sep 1999.

[44] G. R. Cleaves, T. E. Ryan, and R. W. Schlesinger. Identification and characterization of type 2 dengue virus replicative intermediate and replicative form RNAs. Virology, 111(1):73–83, May 1981.

[45] H. Wang and A. W. Tai. Continuous de novo generation of spatially segregated hepatitis C virus replication organelles revealed by pulse-chase imaging. J. Hepatol., 66(1):55–66, 01 2017.

[46] O. Grünvogel, O. Colasanti, J. Y. Lee, V. Klöss, S. Belouzard, A. Reustle, K. Esser-Nobis, J. Hesebeck-Brinckmann, P. Mutz, K. Hoffmann, A. Mehrabi, R. Koschny, F. W. R. Vondran, D. Gotthardt, P. Schnitzler, C. Neumann-Haefelin, R. Thimme, M. Binder, R. Bartenschlager, J. Dubuisson, A. H. Dalpke, and V. Lohmann. Secretion of Hepatitis C Virus Replication Intermediates Reduces Activation of Toll-Like Receptor 3 in Hepatocytes. Gastroenterology, 154(8):2237–2251, 06 2018.

[47] C. Wang, J. Pflugheber, R. Sumpter, D. L. Sodora, D. Hui, G. C. Sen, and M. Gale. Alpha interferon induces distinct translational control programs to suppress hepatitis C virus RNA replication. J. Virol., 77(7):3898–3912, Apr 2003.

[48] Margaret Robinson, Huiling Yang, Siu-Chi Sun, Betty Peng, Yang Tian, Nikos Pagratis, Andrew E Greenstein, and William E Delaney. Novel hepatitis C virus reporter replicon cell lines enable efficient antiviral screening against genotype 1a. Antimicrob. Agents Chemother., 54(8):3099–3106, 2010.

[49] J. J. Arnold and C. E. Cameron. Poliovirus RNA-dependent RNA polymerase (3D(pol)). Assembly of stable, elongation-competent complexes by using a symmetrical primer-template substrate (sym/sub). J. Biol. Chem., 275(8):5329–5336, Feb 2000.

[50] D. Baltimore. Structure of the poliovirus replicative intermediate RNA. J. Mol. Biol., 32(2):359–368, Mar 1968.

[51] O. C. Richards, S. C. Martin, H. G. Jense, and E. Ehrenfeld. Structure of poliovirus replicative intermediate RNA. Electron microscope analysis of RNA cross-linked in vivo with psoralen derivative. J. Mol. Biol., 173(3):325–340, Mar 1984.

[52] L. Liu, H. Dong, H. Chen, J. Zhang, H. Ling, Z. Li, P. Y. Shi, and H. Li. Flavivirus RNA cap methyltransferase: structure, function, and inhibition. Front Biol (Beijing), 5(4):286–303, Aug 2010.

[53] J. Pelletier and N. Sonenberg. Internal initiation of translation of eukaryotic mRNA directed by a sequence derived from poliovirus RNA. Nature, 334(6180):320–325, Jul 1988.

[54] K. Tsukiyama-Kohara, N. Iizuka, M. Kohara, and A. Nomoto. Internal ribosome entry site within hepatitis C virus RNA. J. Virol., 66(3):1476–1483, Mar 1992.

[55] R. J. Kuhn, W. Zhang, M. G. Rossmann, S. V. Pletnev, J. Corver, E. Lenches, C. T. Jones, S. Mukhopadhyay, P. R. Chipman, E. G. Strauss, T. S. Baker, and J. H. Strauss. Structure of dengue virus: implications for flavivirus organization, maturation, and fusion. Cell, 108(5):717–725, Mar 2002.

[56] Christian G Schüttler, Christine Thomas, Thomas Discher, Georg Friese, Jürgen Lohmeyer, Ralph Schuster, Stephan Schaefer, and Wolfram H Gerlich. Variable ratio of hepatitis C virus RNA to viral core antigen in patient sera. J. Clin. Microbiol., 42(5):1977–1981, May 2004.

[57] J. Y. Lin, T. C. Chen, K. F. Weng, S. C. Chang, L. L. Chen, and S. R. Shih. Viral and host proteins involved in picornavirus life cycle. J. Biomed. Sci., 16:103, Nov 2009.

[58] S. J. Keum, S. M. Park, J. H. Park, J. H. Jung, E. J. Shin, and S. K. Jang. The specific infectivity of hepatitis C virus changes through its life cycle. Virology, 433(2):462–470, Nov 2012.

[59] N. V. Ivanisenko, E. L. Mishchenko, I. R. Akberdin, P. S. Demenkov, V. A. Likhoshvai, K. N. Kozlov, D. I. Todorov, V. V. Gursky, M. G. Samsonova, A. M. Samsonov, D. Clausznitzer, L. Kaderali, N. A. Kolchanov, and V. A. Ivanisenko. A new stochastic model for subgenomic hepatitis C virus replication considers drug resistant mutants. PLoS ONE, 9(3):e91502, 2014.

[60] S. Marino, I. B. Hogue, C. J. Ray, and D. E. Kirschner. A methodology for performing global uncertainty and sensitivity analysis in systems biology. J. Theor. Biol., 254(1):178–196, Sep 2008.

[61] Jérôme Gouttenoire, Philippe Roingeard, François Penin, and Darius Moradpour. Amphipathic α-helix AH2 is a major determinant for the oligomerization of hepatitis C virus nonstructural protein 4B. “J. Virol”., 84(24):12529–12537, 2010.

[62] A. Diaz, X. Wang, and P. Ahlquist. Membrane-shaping host reticulon proteins play crucial roles in viral RNA replication compartment formation and function. Proc. Natl. Acad. Sci. U.S.A., 107(37):16291–16296, Sep 2010.

[63] A. Diaz and P. Ahlquist. Role of host reticulon proteins in rearranging membranes for positive-strand RNA virus replication. Curr. Opin. Microbiol., 15(4):519–524, Aug 2012.

[64] K. L. Berger, J. D. Cooper, N. S. Heaton, R. Yoon, T. E. Oakland, T. X. Jordan, G. Mateu, A. Grakoui, and G. Randall. Roles for endocytic trafficking and phosphatidylinositol 4-kinase III alpha in hepatitis C virus replication. Proc. Natl. Acad. Sci. U.S.A., 106(18):7577–7582, May 2009.

[65] K. L. Berger and G. Randall. Potential roles for cellular cofactors in hepatitis C virus replication complex formation. Commun Integr Biol, 2(6):471–473, Nov 2009.

[66] M. Arita, H. Kojima, T. Nagano, T. Okabe, T. Wakita, and H. Shimizu. Phosphatidylinositol 4-kinase III beta is a target of enviroxime-like compounds for antipoliovirus activity. J. Virol., 85(5):2364–2372, Mar 2011.

[67] H. Zhu, F. Wong-Staal, H. Lee, A. Syder, J. McKelvy, R. T. Schooley, and D. L. Wyles. Evaluation of ITX 5061, a scavenger receptor B1 antagonist: resistance selection and activity in combination with other hepatitis C virus antivirals. J. Infect. Dis., 205(4):656–662, Feb 2012.

[68] F. Xiao, I. Fofana, C. Thumann, L. Mailly, R. Alles, E. Robinet, N. Meyer, M. Schaeffer, F. Habersetzer, M. Doffoël, P. Leyssen, J. Neyts, M. B. Zeisel, and T. F. Baumert. Synergy of entry inhibitors with direct-acting antivirals uncovers novel combinations for prevention and treatment of hepatitis C. Gut, 64(3):483–494, Mar 2015.

[69] CI Bliss. The toxicity of poisons applied jointly 1. Annals of applied biology, 26(3):585–615, 1939.

[70] M. Comas-Garcia. Packaging of Genomic RNA in Positive-Sense Single-Stranded RNA Viruses: A Complex Story. Viruses, 11(3), 03 2019.

[71] Y. Song, O. Gorbatsevych, Y. Liu, J. Mugavero, S. H. Shen, C. B. Ward, E. Asare, P. Jiang, A. V. Paul, S. Mueller, and E. Wimmer. Limits of variation, specific infectivity, and genome packaging of massively recoded poliovirus genomes. Proc. Natl. Acad. Sci. U.S.A., 114(41):E8731–E8740, 10 2017.

[72] E. R. Aguilera, A. K. Erickson, P. R. Jesudhasan, C. M. Robinson, and J. K. Pfeiffer. Plaques Formed by Mutagenized Viral Populations Have Elevated Coinfection Frequencies. mBio, 8(2), 03 2017.

[73] J. R. Coleman, D. Papamichail, S. Skiena, B. Futcher, E. Wimmer, and S. Mueller. Virus attenuation by genome-scale changes in codon pair bias. Science, 320(5884):1784–1787, Jun 2008.

[74] V. Lohmann, S. Hoffmann, U. Herian, F. Penin, and R. Bartenschlager. Viral and cellular determinants of hepatitis C virus RNA replication in cell culture. J. Virol., 77(5):3007–3019, Mar 2003.

[75] K. J. Blight, J. A. McKeating, and C. M. Rice. Highly permissive cell lines for subgenomic and genomic hepatitis C virus RNA replication. J. Virol., 76(24):13001–13014, Dec 2002.

[76] Ankit Rohatgi. Webplotdigitizer: Version 4.3, 2020.

[77] M. A. Beaumont, W. Zhang, and D. J. Balding. Approximate Bayesian computation in population genetics. Genetics, 162(4):2025–2035, Dec 2002.

[78] Andrea Saltelli and Ricardo Bolado. An alternative way to compute Fourier amplitude sensitivity test (FAST). Computational Statistics & Data Analysis, 26(4):445–460, 1998.

[79] Daniel T Gillespie. Exact stochastic simulation of coupled chemical reactions. The journal of physical chemistry, 81(25):2340–2361, 1977.

## References

[1] T. R. Aunins, K. A. Marsh, G. Subramanya, S. L. Uprichard, A. S. Perelson, and A. Chatterjee. Intracellular Hepatitis C Virus Modeling Predicts Infection Dynamics and Viral Protein Mechanisms. J. Virol., 92(11), 06 2018.

[2] R. Bartenschlager, M. Frese, and T. Pietschmann. Novel insights into hepatitis C virus replication and persistence. Adv. Virus Res., 63:71–180, 2004.

[3] M. A. Beaumont, W. Zhang, and D. J. Balding. Approximate Bayesian computation in population genetics. Genetics, 162(4):2025–2035, Dec 2002.

[4] Marco Binder, Nurgazy Sulaimanov, Diana Clausznitzer, Manuel Schulze, Christian M Hüber, Simon M Lenz, Johannes P Schlöder, Martin Trippler, Ralf Bartenschlager, Volker Lohmann, et al. Replication vesicles are load-and choke-points in the hepatitis c virus lifecycle. PLoS Pathog., 9(8), 2013.

[5] B. Brandenburg, L. Y. Lee, M. Lakadamyali, M. J. Rust, X. Zhuang, and J. M. Hogle. Imaging poliovirus entry in live cells. PLoS Biol., 5(7):e183, Jul 2007.

[6] Graham R Cleaves and Donald T Dubin. Methylation status of intracellular dengue type 2 40 s RNA. Virology, 96(1):159–165, 1979.

[7] M. Comas-Garcia. Packaging of Genomic RNA in Positive-Sense Single-Stranded RNA Viruses: A Complex Story. Viruses, 11(3), 03 2019.

[8] M. Cortese, S. Goellner, E. G. Acosta, C. J. Neufeldt, O. Oleksiuk, M. Lampe, U. Haselmann, C. Funaya, N. Schieber, P. Ronchi, M. Schorb, P. Pruunsild, Y. Schwab, L. Chatel-Chaix, A. Ruggieri, and R. Bartenschlager. Ultrastructural Characterization of Zika Virus Replication Factories. Cell Rep, 18(9):2113–2123, 02 2017.

[9] H. Dahari, R. M. Ribeiro, C. M. Rice, and A. S. Perelson. Mathematical Modeling of Subgenomic Hepatitis C Virus Replication in Huh-7 Cells. J. Virol., 81(2):750–760, Jan 2007.

[10] J. A. den Boon and P. Ahlquist. Organelle-like membrane compartmentalization of positive-strand RNA virus replication factories. Annu. Rev. Microbiol., 64:241–256, 2010.

[11] Arturo Diaz, Xiaofeng Wang, and Paul Ahlquist. Membrane-shaping host reticulon proteins play crucial roles in viral RNA replication compartment formation and function. Proceedings of the National Academy of Sciences, 107(37):16291–16296, 2010.

[12] O. Grünvogel, O. Colasanti, J. Y. Lee, V. Klöss, S. Belouzard, A. Reustle, K. Esser-Nobis, J. Hesebeck- Brinckmann, P. Mutz, K. Hoffmann, A. Mehrabi, R. Koschny, F. W. R. Vondran, D. Gotthardt, P. Schnitzler, C. Neumann-Haefelin, R. Thimme, M. Binder, R. Bartenschlager, J. Dubuisson, A. H. Dalpke, and V. Lohmann. Secretion of Hepatitis C Virus Replication Intermediates Reduces Activation of Toll-Like Receptor 3 in Hepatocytes. Gastroenterology, 154(8):2237–2251, 06 2018.

[13] Nikita V Ivanisenko, Elena L Mishchenko, Ilya R Akberdin, Pavel S Demenkov, Vitaly A Likhoshvai, Konstantin N Kozlov, Dmitry I Todorov, Vitaly V Gursky, Maria G Samsonova, Alexander M Samsonov, et al. A new stochastic model for subgenomic hepatitis c virus replication considers drug resistant mutants. PloS one, 9(3):e91502, 2014.

[14] T. Kazakov, F. Yang, H. N. Ramanathan, A. Kohlway, M. S. Diamond, and B. D. Lindenbach. Hepatitis C virus RNA replication depends on specific cis- and trans-acting activities of viral nonstructural proteins. PLoS Pathog., 11(4):e1004817, Apr 2015.

[15] S. J. Keum, S. M. Park, J. H. Park, J. H. Jung, E. J. Shin, and S. K. Jang. The specific infectivity of hepatitis C virus changes through its life cycle. Virology, 433(2):462–470, Nov 2012.

[16] Kévin Knoops, Montserrat Bárcena, Ronald WAL Limpens, Abraham J Koster, A Mieke Mommaas, and Eric J Snijder. Ultrastructural characterization of arterivirus replication structures: Reshaping the endoplasmic reticulum to accommodate viral RNA synthesis. J. Virol., 86(5):2474–2487, Mar 2012.

[17] R. J. Kuhn, W. Zhang, M. G. Rossmann, S. V. Pletnev, J. Corver, E. Lenches, C. T. Jones, S. Mukhopadhyay, P. R. Chipman, E. G. Strauss, T. S. Baker, and J. H. Strauss. Structure of dengue virus: implications for flavivirus organization, maturation, and fusion. Cell, 108(5):717–725, Mar 2002.

[18] Q. Li, Y. Tong, Y. Xu, J. Niu, and J. Zhong. Genetic Analysis of Serum-Derived Defective Hepatitis C Virus Genomes Revealed Novel Viral cis Elements for Virus Replication and Assembly. J. Virol., 92(7), 04 2018.

[19] Ronald WAL Limpens, Hilde M van der Schaar, Darshan Kumar, Abraham J Koster, Eric J Snijder, Frank JM van Kuppeveld, and Montserrat Bárcena. The transformation of enterovirus replication structures: a three-dimensional study of single-and double-membrane compartments. MBio, 2(5):e00166–11, 2011.

[20] J. Y. Lin, T. C. Chen, K. F. Weng, S. C. Chang, L. L. Chen, and S. R. Shih. Viral and host proteins involved in picornavirus life cycle. J. Biomed. Sci., 16:103, Nov 2009.

[21] Lihui Liu, Hongping Dong, Hui Chen, Jing Zhang, Hua Ling, Zhong Li, Pei-Yong Shi, and Hongmin Li. Flavivirus RNA cap methyltransferase: structure, function, and inhibition. Frontiers in biology, 5(4):286–303, 2010.

[22] Michael D McKay, Richard J Beckman, and William J Conover. Comparison of three methods for selecting values of input variables in the analysis of output from a computer code. Technometrics, 21(2):239–245, 1979.

[23] Alison K McLean, Fabio Luciani, and Mark M Tanaka. Trade-offs in resource allocation in the intracellular life-cycle of hepatitis c virus. Journal of theoretical biology, 267(4):565–572, 2010.

[24] Sven Miller and Jacomine Krijnse-Locker. Modification of intracellular membrane structures for virus replication. Nature Reviews Microbiology, 6(5):363–374, 2008.

[25] Jerry Pelletier and Nahum Sonenberg. Internal initiation of translation of eukaryotic mRNA directed by a sequence derived from poliovirus RNA. Nature, 334(6180):320–325, 1988.

[26] I. Romero-Brey and R. Bartenschlager. Membranous replication factories induced by plus-strand RNA viruses. Viruses, 6(7):2826–2857, Jul 2014.

[27] I. Romero-Brey, A. Merz, A. Chiramel, J. Y. Lee, P. Chlanda, U. Haselman, R. Santarella-Mellwig, A. Habermann, S. Hoppe, S. Kallis, P. Walther, C. Antony, J. Krijnse-Locker, and R. Bartenschlager. Three-dimensional architecture and biogenesis of membrane structures associated with hepatitis c virus replication. PLoS Pathog., 8(12):e1003056, 2012.

[28] M. B. Schulte, J. A. Draghi, J. B. Plotkin, and R. Andino. Experimentally guided models reveal replication principles that shape the mutation distribution of RNA viruses. Elife, 4, Jan 2015.

[29] Christian G Schüttler, Christine Thomas, Thomas Discher, Georg Friese, Jürgen Lohmeyer, Ralph Schuster, Stephan Schaefer, and Wolfram H Gerlich. Variable ratio of hepatitis c virus RNA to viral core antigen in patient sera. J. Clin. Microbiol., 42(5):1977–1981, May 2004.

[30] Y. Song, O. Gorbatsevych, Y. Liu, J. Mugavero, S. H. Shen, C. B. Ward, E. Asare, P. Jiang, A. V. Paul, S. Mueller, and E. Wimmer. Limits of variation, specific infectivity, and genome packaging of massively recoded poliovirus genomes. Proc. Natl. Acad. Sci. U.S.A., 114(41):E8731–E8740, 10 2017.

[31] Kyoko Tsukiyama-Kohara, N Iizuka, M Kohara, and A Nomoto. Internal ribosome entry site within hepatitis C virus RNA. Journal of virology, 66(3):1476–1483, 1992.

[33] M. Ulasli, M. H. Verheije, C. A. de Haan, and F. Reggiori. Qualitative and quantitative ultrastructural analysis of the membrane rearrangements induced by coronavirus. Cell. Microbiol., 12(6):844–861, Jun 2010.

[34] Barbara van der Hoeven, Diede Oudshoorn, Abraham J Koster, Eric J Snijder, Marjolein Kikkert, and Montserrat Bárcena. Biogenesis and architecture of arterivirus replication organelles. Virus research, 220:70–90, 2016.

[35] H. Wang and A. W. Tai. Continuous de novo generation of spatially segregated hepatitis C virus replication organelles revealed by pulse-chase imaging. J. Hepatol., 66(1):55–66, 01 2017.

[36] Ming-Jhan Wu, Po-Yuan Ke, John T-A Hsu, Chau-Ting Yeh, and Jim-Tong Horng. Reticulon 3 interacts with ns4b of the hepatitis c virus and negatively regulates viral replication by disrupting ns4b self-interaction. Cellular microbiology, 16(11):1603–1618, 2014.

[37] Carolin Zitzmann, Bianca Schmid, Alessia Ruggieri, Alan S Perelson, Marco Binder, Ralf Bartenschlager, and Lars Kaderali. A coupled mathematical model of the intracellular replication of dengue virus and the host cell immune response to infection. Frontiers in microbiology, 11:725, 2020.

